# High-resolution molecular atlas of a lung tumor in 3D

**DOI:** 10.1101/2023.05.10.539644

**Authors:** Tancredi Massimo Pentimalli, Simon Schallenberg, Daniel León-Periñán, Ivano Legnini, Ilan Theurillat, Gwendolin Thomas, Anastasiya Boltengagen, Sonja Fritzsche, Jose Nimo, Lukas Ruff, Gabriel Dernbach, Philipp Jurmeister, Sarah Murphy, Mark T. Gregory, Yan Liang, Michelangelo Cordenonsi, Stefano Piccolo, Fabian Coscia, Andrew Woehler, Nikos Karaiskos, Frederick Klauschen, Nikolaus Rajewsky

## Abstract

Cells live and interact in three-dimensional (3D) cellular neighborhoods. However, histology and spatial omics methods mostly focus on 2D tissue sections. Here we present a 3D spatial atlas of a routine clinical sample, an aggressive human lung carcinoma, by combining *in situ* quantification of 960 cancer-related genes across ∼340,000 cells with measurements of tissue-mechanical components. 3D cellular neighborhoods subdivided the tumor microenvironment into tumor, stromal, and immune multicellular niches. Interestingly, pseudotime analysis suggested that pro-invasive epithelial-to-mesenchymal transition (EMT), detected in stroma-infiltrating tumor cells, already occurred in one region at the tumor surface. There, myofibroblasts and macrophages specifically co-localized with pre-invasive tumor cells and their multicellular molecular signature identified patients with shorter survival. Moreover, cytotoxic T-cells did not infiltrate this niche but colocalized with inhibitory dendritic and regulatory T cells. Importantly, systematic scoring of cell-cell interactions in 3D neighborhoods highlighted niche-specific signaling networks accompanying tumor invasion and immune escape. Compared to 2D, 3D neighborhoods improved the characterization of immune niches by identifying dendritic niches, capturing the 3D extension of T-cell niches and boosting the quantification of niche-specific cell-cell interactions, including druggable immune checkpoints. We believe that 3D communication analyses can improve the design of clinical studies investigating personalized, combination immuno-oncology therapies.

## INTRODUCTION

Cells build three-dimensional (3D) tissues to perform specific functions. Mechanical cues and molecular signaling regulating tissue development and function operate in 3D between neighboring cells. The comprehensive characterization of 3D cellular neighborhoods in terms of cellular composition, molecular states, mechanical properties and ligand-receptor interactions is thus central to understand tissue function in health and disease. While light microscopy [1] and hematoxylin and eosin (H&E) staining [2] enabled the morphological characterization of tissue organization in 3D, the molecular exploration of 3D cellular neighborhoods is still in its infancy.

Recent developments in spatial omics methods profoundly changed our ability to characterize tissue molecular properties [3]. While STARmap [4] and DISCO-MS [5] enabled the 3D profiling of intact mouse tissues, *ad hoc* tissue processing hinders their application to routinely collected fresh-frozen (FF) and formalin-fixed paraffin-embedded (FFPE) clinical samples. Therefore, the initial molecular studies leveraged the computational registration of sequential 2D sections for the *in silico* reconstruction of tissue architecture in 3D. These included the imaging mass cytometry (IMC) study of breast cancer (40 proteins in 3 samples) [6] and the multiplexed immunofluorescence (IF) study of colorectal cancer (32 proteins in 1 sample) [7]. While enabling single cell-resolved tissue profiling, these methods are limited in plexity, only measuring tens of proteins at the time. Conversely, sequencing-based spatial transcriptomics methods, such as Slide-seq [8], ST [9] (now Visium) and Stereo-seq [10], which rely on the capture of polyadenylated transcripts on spatially barcoded regions, enabled the unbiased, genome-wide characterization of 3D gene expression patterns in the mouse hippocampus (10 µm beads) [8], rheumatoid arthritis synovium (100 µm spots) [11] and developing Drosophila embryo (10 µm x 10µm or 25 µm x25 µm bins) [12]. However, intrinsic resolution limits and difficulties in assigning regions to single cells prevented single-cell resolved analyses. Furthermore, polyA-based methods require intact RNA molecules (as in FF tissues) for efficient transcript capture, thus preventing their application to fragmented RNA typical of FFPE clinical samples. On the other hand, high-plex imaging-based spatial transcriptomic methods, such as MERFISH [13] and CosMx [14], use probes to recognize their targets with single molecule resolution, are compatible with both FF and FFPE tissues but have not yet been used for the 3D reconstruction of clinical specimens.

Here we present the 3D, high-plex and single-cell resolved molecular atlas of a human tissue, enabling the reconstruction, multimodal profiling and systematic exploration of 3D cellular neighborhoods using CosMx. For this proof-of-principle study, we focused on an aggressive, early-stage non-small cell lung cancer (NSCLC) sample and investigated the ability of 3D neighborhoods to identify tumor invasion and immune escape mechanisms active in the patient under study. Despite being a paradigm of precision medicine with an arsenal of molecularly targeted therapies and immunotherapy agents [15], NSCLC remains the main cause of cancer-related death [16]. Furthermore, only 20% of NSCLC patients respond to immunotherapy [17] and up to 30% of treated patients suffer from immune-related adverse events [18]. Therefore, the identification of personalized immunotherapy biomarkers represents an urgent unmet clinical need to maximize therapeutic efficacy and limit toxicity in NSCLC.

As DNA sequencing guides the administration of targeted therapies through the detection of druggable mutations [19], spatial omics methods hold great promise for the identification of immune escape mechanisms active in individual patients, which could serve as personalized (immune)therapy biomarkers. Here we showcase the power of high-plex, single cell resolved, molecular histology to simultaneously profile cell types, molecular states, and receptor-ligand interactions underlying tumor invasion and immune escape. Importantly, we demonstrate how 3D neighborhoods improved the identification of tumor immune escape mechanisms, compared to their 2D counterparts.

## RESULTS

### 1. Multimodal study of one aggressive NSCLC tumor in 3D

For this proof-of-principle study, we selected an early-stage NSCLC patient, which demonstrated rapid disease progression and aggressive tumor biology (**Ext data fig 1a**). To reconstruct 3D cellular neighborhoods, we cut 34 consecutive 5 µm-thick tissue sections from a routinely collected, archival FFPE tumor block. We leveraged deep-learning based classification of tumor, stromal and normal lung-resident compartments in one whole-slide H&E image to select one 16 mm^2^ wide region of interest (ROI) featuring the copresence of both the primary tumor and cancer-associated stroma for spatial transcriptomics investigation (**Ext data fig 1b**). Furthermore, we reasoned that the presence of small-caliber airways crossing the section planes would enable the assessment of section alignments in 3D. We then collected single-cell resolved spatial transcriptomics data from every 6^th^ section (30µm section-to-section interval) and profiled intervening sections with complementary modalities, enabling the integrative analysis of tissue morphology, ECM composition, protein markers and gene expression (**Fig 1a**).

**Figure 1.**
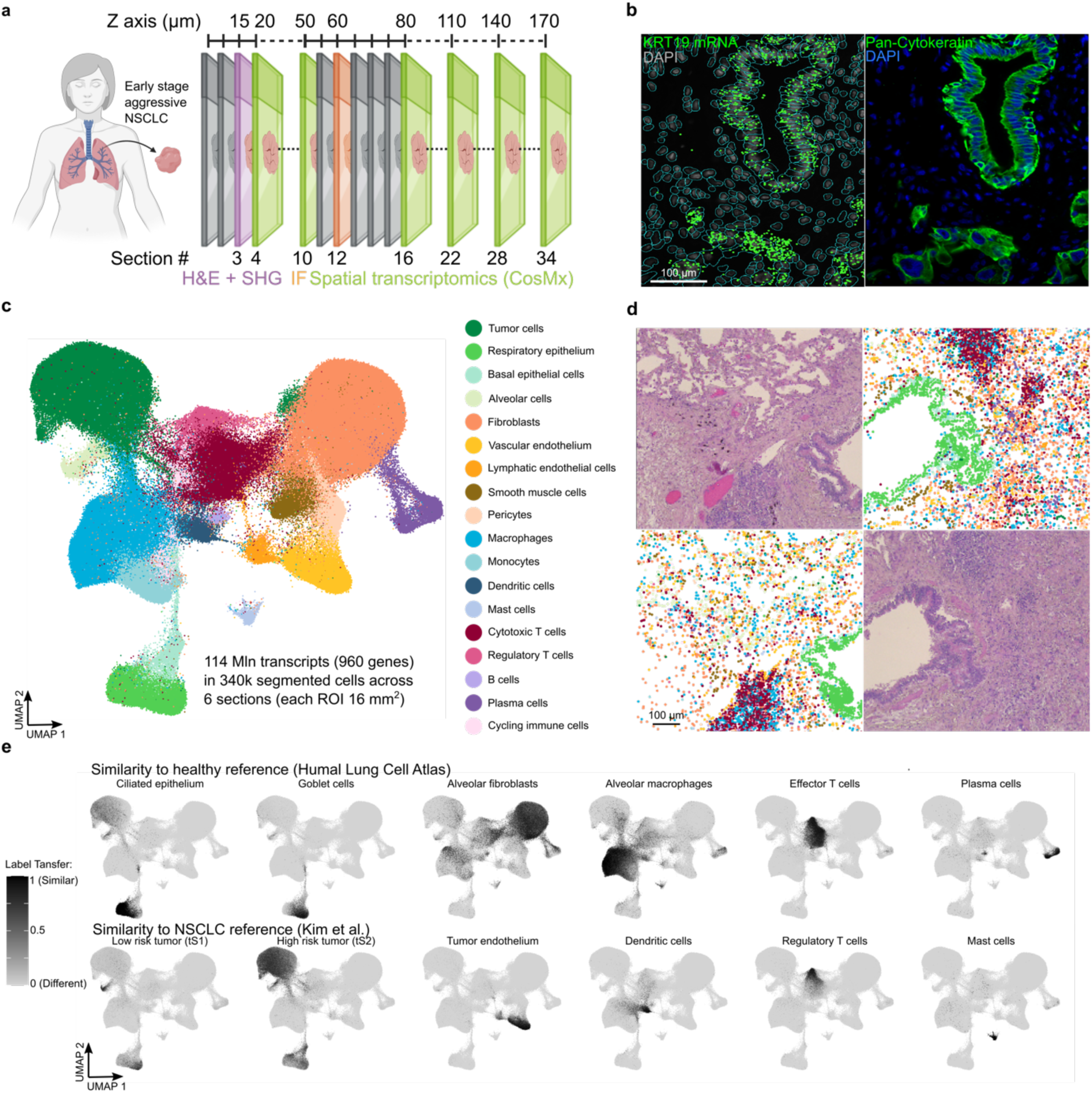
Molecular histology of the tumor microenvironment at single cell resolution. **a)** Experimental design for the 3D reconstruction of cellular neighborhoods. 34 consecutive 5 µm sections were cut from one non-small cell lung cancer (NSCLC) tumor block. Second harmonic imaging (SHG, quantifies collagen and elastin), hematoxylin and eosin (H&E, detects tissue morphology), spatial transcriptomics (1000-plex RNA in situ hybridization with CosMx Spatial Molecular Imager), immunofluorescence (IF) were combined for multimodal spatial profiling. **b)** Keratin 19 (KRT19) mRNA localization matches pan-cytokeratin immunostaining. Left: *KRT19* mRN A captured by CosMx (green: *KRT19* transcripts, gray: DAPI (nuclei), blue: cell segmentation masks). Right: pan-cytokeratin IF (epithelial cells, green; DAPI: blue) **c)** UMAP of cell types in the tumor microenvironment. **d)** Traditional and molecular histology. Top left and bottom right: H&E staining (section 3). Top right and bottom left: Cell types mapped to their spatial position (section 4). **e)** Congruence of gene expression profiles in segmented cells with healthy and tumor single cell RNA sequencing references. UMAP plots are colored by their Label Transfer scores indicating similarity to reference cell types.

### 2. Molecular histology of the tumor microenvironment at single-cell resolution

To gain insights into the spatial organization and crosstalk between tumor, stromal and immune populations, we performed 1000-plex RNA in situ hybridization (ISH) with CosMx Spatial Molecular Imager and leveraged deep-learning based cell segmentation of IF staining of nuclei and plasma membranes to obtain single cell gene expression profiles (Methods). We profiled 960 cancer-related genes allowing the simultaneous characterization of cellular identities (211 genes), transcriptomic states (268 genes), and cell-to-cell communication events (517 genes) in the TME [14] (**Supplementary table 1**).

We imaged a total of 155,055,865 transcripts distributed in six sequential, non-consecutive ROIs (**Ext data fig 1c**). 20 negative probes (i.e. targeting sequences not present in human tissues) composed only 0.27% of the total detected molecules, demonstrating highly specific *in-situ* detection of target genes. To further assess data quality, we confirmed the colocalization of pancytokeratin (panCK) staining and ISH signal for *KRT19* transcripts in the same section (**Fig 1b**). 74.1% of the detected transcripts were assigned to 340,644 segmented cells (101 median genes and 198 median transcripts/cell). Near-perfect correlations of total transcript counts confirmed robust gene expression profiling across sections (**Ext data fig 1d**). Furthermore, hierarchical clustering recapitulated the spatial arrangement as sections closer in the z-axis showed slightly higher correlations than distant ones.

Through unsupervised clustering of gene expression in segmented cells (**Ext data fig 1e**), we identified 18 cell types (**Fig 1c**) based on the expression of canonical marker genes (**Ext data fig 1f**) and IF positivity to panCK staining (**Ext data fig 1g**). In line with previous reports [20], tumor cells were characterized by a larger cell size and higher transcript counts (**Ext data fig 1g**). Transferring cell type annotations back to their tissue positions, we observed their correspondence with the spatial patterns of epithelial, stromal and immune populations observed using routine histology (**Fig 1d, Ext data fig 1h-i**). Comparing gene expression of segmented cells with published single-cell RNA sequencing atlases, we confirmed the agreement of our cell type annotations with healthy lung [21] and NSCLC atlases [22] (**Fig 1e**).

In summary, we generated a high-quality, single cell-resolved molecular atlas of the TME, including more than 340,000 cells spanning 18 epithelial, stromal and immune cell types and went on to explore their 3D cellular neighborhoods.

### 3. Reconstruction of 3D cellular neighborhoods

To reconstruct 3D cellular neighborhoods, we leveraged STIM [23], which employs state-of-the-art computer vision techniques, to computationally align spatial transcriptomics data and generate a 3D molecular map of the TME at single-cell resolution. Supporting the high fidelity of our 3D model, visualization of respiratory epithelium cells revealed that the successful reconstruction of airway lumen moving along the individual 2D planes (**Fig 2a**). We thus went on to define 3D neighborhoods for each cell as a spherical space encompassing all cells located within a 50 µm center-to-center distance (**Fig 2b**). By design, 3D cellular neighborhoods included both a circular 314.16 µm^2^ area in the section where the center cell is located and two 251.33 µm^2^ areas in the sections immediately above and below in the z plane, which represents a 2.6-fold increase compared to the study of 2D cellular neighborhoods.

**Figure 2.**
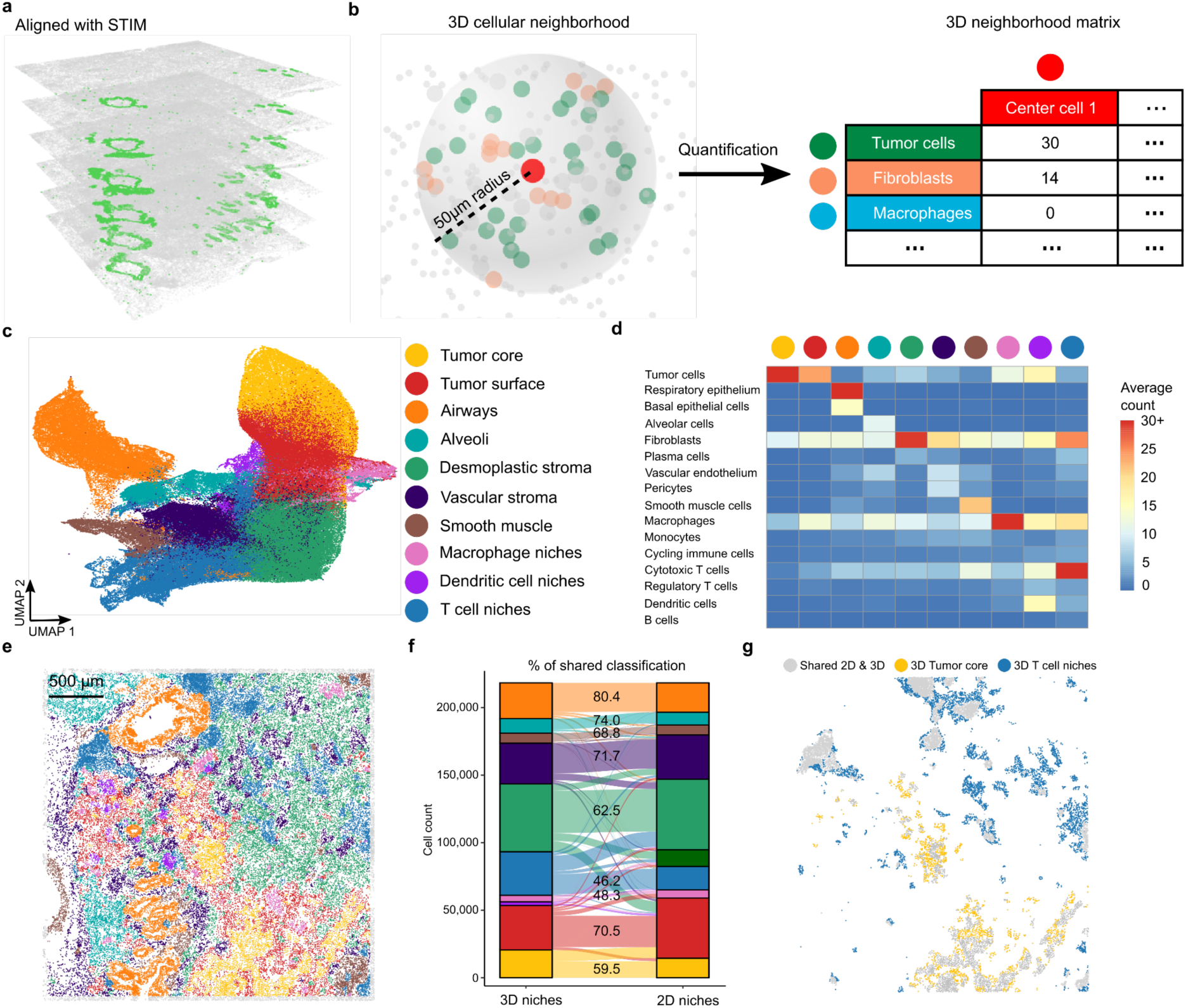
3D neighborhoods identify multicellular niches in the tumor microenvironment. **a)** 3D plot of cells in the 6 CosMx sections after registration. Green: respiratory epithelium cells, gray: other cell types. Axes are scaled to the same length. **b)** Analysis of 3D cellular neighborhoods. Exemplary 3D neighborhood (left). Red: center cell: red, dark green: tumor cells, orange: fibroblasts, gray: other cell types. Quantification of cell types in 3D neighborhoods is used to build the neighborhood matrix (right). Tumor cells, fibroblasts and macrophages are shown out of 18 cell types quantified. **c)** UMAP of 3D cellular neighborhoods. Cells are grouped based on their 3D neighborhood composition, regardless of their gene expression, and colored by 3D niche assignment. **d)** Composition of 3D neighborhoods in multicellular niches. Heatmap of niche-specific average cell type 3D neighborhood counts. Color scale is clipped to 30 for visualization purposes. **e)** Spatial map of 3D multicellular niches (section 10). Color legend in panel c. gray: cells within 50 µm of the section edge. **f)** Comparison of 2D and 3D niches. Alluvial plot of niche assignments when analyzing 2D and 3D neighborhoods. Lines follow the assignment of the same cell in 3D (left) and 2D niches (left). Color legend in panel c, dark green: ‘macrophage-rich stroma’. **g)** 3D neighborhoods capture niche spatial continuity. Cells assigned to ‘T cell niches’ and ‘tumor core’ only in 3D (highlighted) connect seemingly separated 2D niches.

Owing to the precise positioning of the ROIs during data collection, relatively minor transformations were required for 3D image registration (**Ext data fig 2a**). Nevertheless, the median shift of cell positions after registration was ∼42 µm, which would have altered 3D cellular neighborhoods if uncorrected.

After excluding cells whose neighborhoods extended beyond the boundaries of the 3D reconstruction (i.e., located in the first/last sections or within 50 µm from the edge of the central sections), we set out to study the 3D neighborhoods for 218,378 center cells.

Quantification of 3D neighborhood cell type composition (**Fig 2b**) revealed the complexity and richness of cellular microenvironments (median of 71 cells from 9 cell types/neighborhood). In comparison, 2D neighborhoods featured not only a reduced number of neighbors but also a lower cell type richness (median of 32 cells from 7 cell types/neighborhood) and a lower alpha diversity – a common measure of species richness in ecology studies – (median Chao index 3D: 10.5 vs 2D: 8) (**Ext data fig 2b**).

### 4. 3D neighborhoods enable the unbiased identification of multicellular niches in the TME

Cellular activities and molecular profiles are shaped by the surrounding tissue microenvironment [3]. To group cells sharing the same tissue microenvironment, we unbiasedly clustered center cells based on their 3D neighborhood cellular composition - regardless of their gene expression-. In this way, we identified 10 TME niches (**Fig 2c**), defined as multicellular compartments sharing a specific neighborhood composition (**Fig 2d**). These comprised both lung-resident epithelial (i.e. ‘airways’ and ‘alveoli’) and stromal (i.e. ‘smooth muscle’) niches and TME-specific tumor, stromal and immune niches. High numbers of tumor cells characterized the 3D neighborhoods of cells assigned to both ‘tumor core’ and the ‘tumor surface’. The tumor core featured a higher cellular density but a lower cellular diversity (**Ext data fig2c**) as 3D neighborhoods were dominated by tumor cells. On the other hand, the tumor surface featured a higher count of fibroblasts, macrophages and cytotoxic T cells, resulting in a higher cell type diversity compatible with its position at the boundary with the surrounding stroma. 3D neighborhoods not only distinguished the ‘vascular stroma’ – rich in vascular endothelium and pericytes – from the ‘desmoplastic stroma’, featuring the highest fibroblast and plasma cell density, but also identified 3 distinct immune niches. Interestingly, tumor cells were abundant in 3D neighborhoods of ‘dendritic cell niches’ and ‘macrophages niches’ but not in ‘T cell niches’. Mapping niche annotations back to their tissue positions revealed that T cell niches were mainly found in proximity of larger airways or dispersed in the stroma. On the other hand, macrophage niches and dendritic cell niches were embedded in the tumor surface (**Fig 2e**), thus explaining the abundance of tumor cells in these niches and suggesting their functional role in the local regulation of anti-tumoral immune responses.

Furthermore, comparison with manual annotations of routine H&E histology by an experienced pathologist (S.S.) confirmed the ability of 3D neighborhoods to capture the TME spatial organization (**Ext data fig2d**).

We then sought to evaluate the ability of 3D neighborhoods to capture the TME spatial organization compared to their 2D counterparts. While 2D niches largely corresponded to 3D ones and even included a ‘macrophage-rich stroma’ (**Ext data fig2e**), 2D neighborhoods did not highlight ‘dendritic niches’. While the 2D and 3D niche assignments agreed for 64.1% of cells, concordance rates varied across niches (**Fig 2f**). In particular, ‘tumor core’ and ‘T cell niches’ showed below average concordance rates as around 50% of cells assigned to these niches in 3D were reassigned in 2D to the ‘desmoplastic stroma’ and ‘tumor surface’, respectively. Consequently, the abundance of ‘T cell niches’ and ‘tumor core’ was reduced in 2D compared to 3D. Importantly, ‘T cell niches’ and ‘tumor core’ formed smaller disconnected patches in 2D and most of the cells unique to the 3D analysis formed bridges across patches, restoring the spatial continuity of these niches (**Fig 2g**).

In summary, we generated a 3D molecular atlas of the TME and leveraged it to study 3D neighborhoods for over 200,000 cells. Remarkably, the unbiased analysis of cell type abundance in 3D neighborhoods decomposed the TME into 10 distinct multicellular niches, corresponding to expert annotations of morphological tissue structures. While 2D niches largely agreed with their 3D counterparts, 3D neighborhoods improved the identification of immune niches in the TME and better captured their 3D spatial continuity.

### 5. Tumor cells acquire pro-invasive molecular programs in the EMT niche

The ability of cancer cells to detach from the epithelial sheet and infiltrate the adjacent stroma (i.e. “invasiveness”) is a hallmark of malignant tumors and essential for metastatic dissemination [24]. At the molecular level, the downregulation of cell-to-cell adhesion molecules (i.e. *EPCAM* and *CDH1*) and the upregulation of mesenchymal markers (i.e. *VIM*) during epithelial-to-mesenchymal transition (EMT) promotes the invasion of surrounding tissues [25]. Quantifying the niche assignments of 38,804 tumor cells, we noted that, while most tumor cells were located in the tumor bed, more than 9,000 tumor cells were assigned to other tissue niches (**Ext data fig3a**). Comparing the observed and expected number of infiltrating tumor cells (Methods), we identified their preferential infiltration of ‘macrophage niches’, ‘dendritic cell niches’, ‘desmoplastic stroma’ and ‘alveolar spaces’ (**Ext data fig3b**). Through 3D rendering, we demonstrate how the ‘tumor surface’ precisely interlocked and covered the ‘tumor core’ and studied the spatial distribution of infiltrating tumor cells (**Fig 3a**). While the spatial continuity of the ‘tumor surface’ with ‘macrophage niches’, ‘dendritic cell niches’ and ‘alveoli’ could explain the enrichment of tumor cells in these niches (**Supp Video 1**), tumor cells the ‘desmoplastic stroma’ extended away from the tumor surface (**Fig 3b**). We thus focused on these cells to identify molecular pathways activated upon tumor invasion.

**Figure 3.**
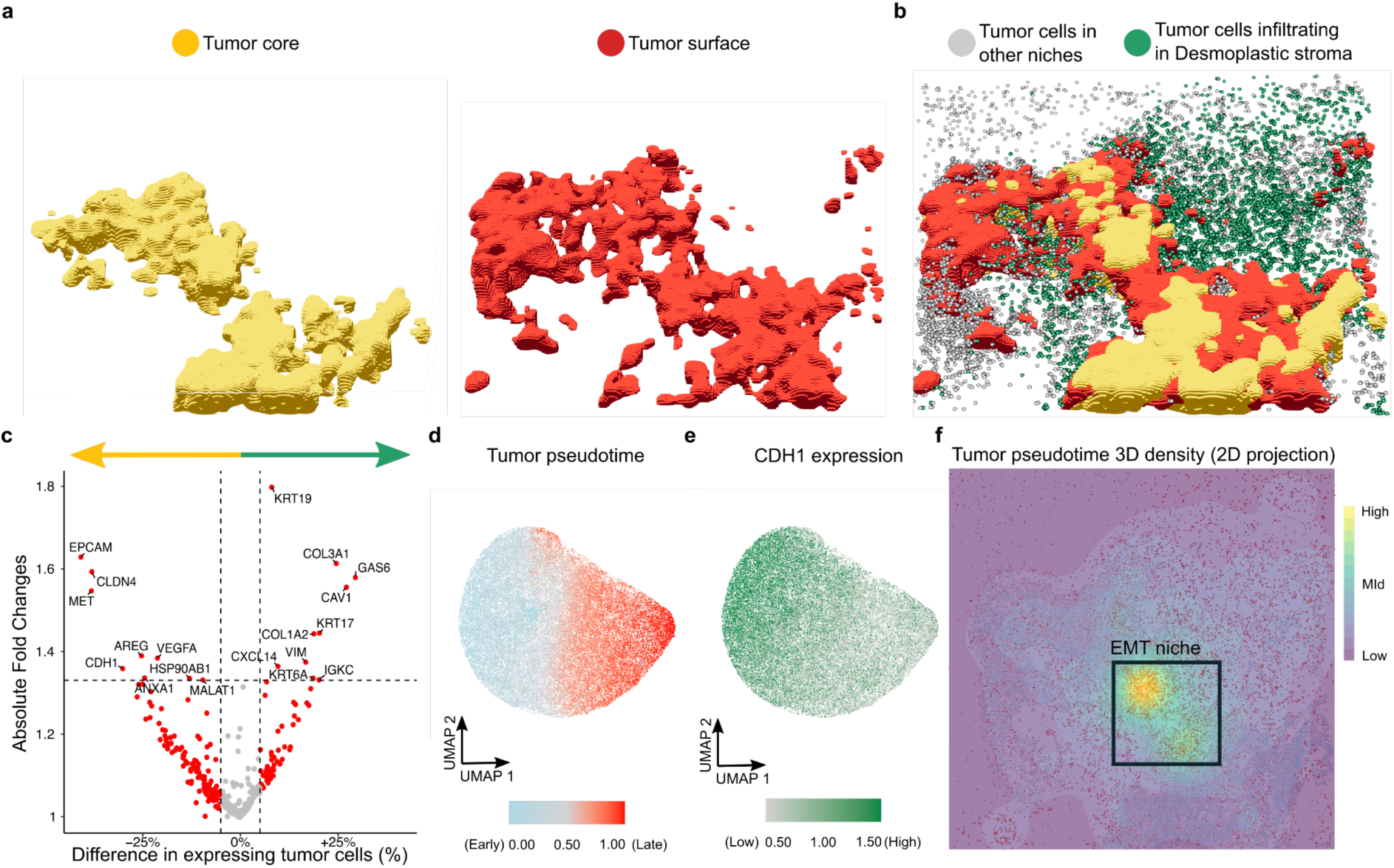
Tumor cells acquire pro-invasive molecular programs in the EMT niche. **a)** 3D rendering of the tumor bed. **b)** 3D rendering of infiltrating tumor cells extending beyond the tumor surface. **c)** Molecular characterization of infiltrating tumor cells. Comparison of gene expression between tumor cells located in the desmoplastic stroma vs those in the tumor core. Red: genes with more > 5% expression difference, labeled: red genes with absolute fold change > 1.33. **d)** Pseudotime captures tumor cell molecular dynamics. UMAP plot of tumor cells based on their transcriptomic profiles. Tumor cells are colored by pseudotime rank. **e)** Epithelial state marker *CDH1* (E-Cadherin) is downregulated with increasing pseudotime. UMAP plot of tumor cells colored by normalized CDH1 expression. **f)** Tumor cells undergo epithelial-to-mesenchymal transition (EMT) in one EMT niche. 3D spatial density of tumor cells with late pseudotime (rank >0.75) highlights one region of the tumor bed.

Comparing the transcriptomic profiles of tumor cells in the ‘desmoplastic stroma’ against those in the ‘tumor core’, we detected the upregulation of cytokeratins (*KRT19*, *KRT17* and *KRT6A*) (**Fig 3c**), which was validated at the protein level by a stronger panCK IF signal (**Ext data fig3c**). Consistent with EMT, *VIM* expression was upregulated in infiltrating tumor cells while *EPCAM* and *CDH1* were downregulated. Interestingly, tumor cells assigned to the ‘tumor surface’ already demonstrated such molecular changes - albeit to a lesser extent - (**Ext data fig 3d**). This is consistent with the view that EMT is a complex, continuous process with a spectrum of intermediate states between a fully epithelial and a fully mesenchymal phenotype [26]. Therefore, we leveraged pseudotime, a popular approach in the single cell transcriptomics field to reconstruct dynamic molecular processes, to rank tumor cells from 0 (early) to 1 (late) (**Fig 3d**). Pseudotime captured EMT dynamics as tumor cells with higher epithelial markers (i.e. *CDH1^high^ EPCAM^high^ VIM^low^*) showed earlier pseudotime, while pseudotime increased in cells undergoing EMT (i.e. *CDH1^low^ EPCAM^low^ VIM^high^*) (**Fig 3e, Ext data fig 3e-f**). Combining tumor pseudotime and 3D niche assignments, we demonstrate the progressive activation of an EMT gene expression program from tumor core, over tumor surface, to stroma infiltrating tumor cells (median pseudotime scores: 0.39, 0.50 and 0.85) (**Ext data fig 3g).** We then evaluated the 3D spatial density of pseudotime scores and confirmed that infiltrating tumor cells feature both an advanced pseudotime and ‘pseudospace’ (intended as distance from the tumor bed), while tumor cells in the tumor core by both an early pseudotime and pseudospace (given their location in the tumor bed).

Strikingly, we discovered one regionat the interface between the ‘tumor surface’ and the ‘desmoplastic stroma’ where the density of mesenchymal-like tumor cells peaked (**Fig 3f**, **Ext data fig 3h**) and IF validated the presence of panCK+ CDH1^low^ cells (**Ext data fig 3i**).

Intriguingly, the EMT niche was characterized by the discrepancy between pseudotime (advanced) and pseudospace (early), indicating that tumor cells may undergo EMT already in the tumor bed before invading the desmoplastic stroma.

### 6. A multicellular molecular signature links the EMT niche with reduced survival in a large cohort

Molecular understanding of tumor invasion is critical for identifying biomarkers to accurately predict and intercept tumor progression [27]. Investigating tumor-intrinsic molecular programs active in the ‘EMT niche’ (**Fig 4a**), we identified *LGALS1* and *NDRG1* as the genes with the strongest differential expression (**Fig 4b**). *LGALS1* (Galectin-1) is known to be upregulated in NSCLC compared to healthy lung tissues [28] and to promote tumor cell EMT and immune escape [29], while *NDRG1* (N-myc downstream-regulated gene 1) promotes stem-like properties in NSCLC cells [30]. Remarkably, their expression was not only enriched but almost spatially restricted to tumor cells in the ‘EMT niche’ (**Fig 4c**, **Ext data fig 4a**).

**Figure 4.**
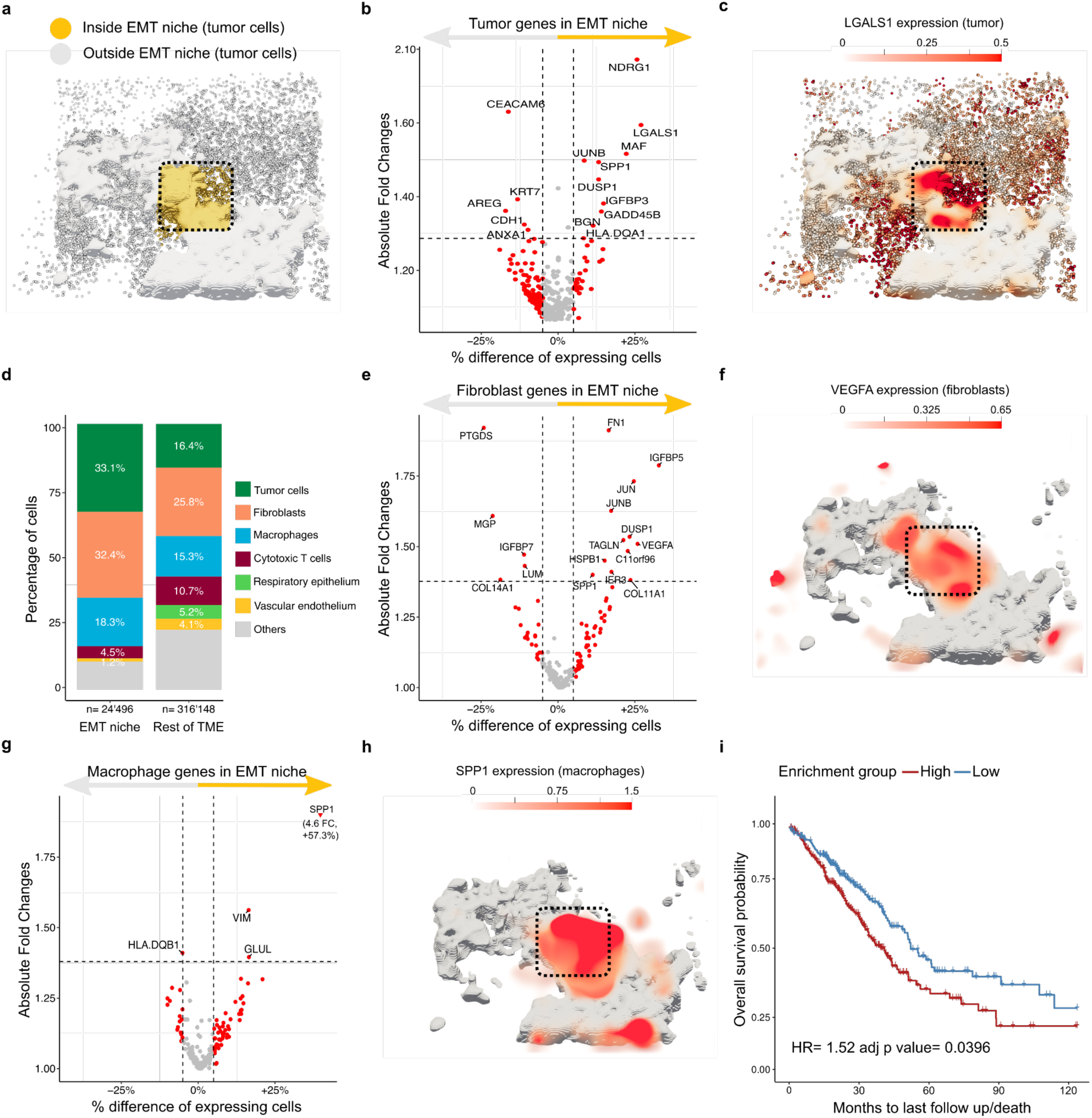
A multicellular molecular signature links the EMT niche with reduced survival in a large cohort. **a)** 3D rendering of EMT niche (yellow). **b)** EMT niche tumor biomarkers. Differential gene expression between tumor cells located inside the EMT niche vs those outside. Red: genes with more > 5% expression difference, labeled: red genes with absolute fold change > 1.33. **c)** *LGALS1* expression is specific for tumor cells in the EMT niche (3D surface rendering), black box: EMT niche. **d)** Tumor cells, fibroblasts and macrophages are the main cell types in the EMT niche. Barplot of cell type composition inside and outside the EMT niche. Gray: cell types with <4% abundance in both regions. **e)** EMT niche fibroblast biomarkers. Differential gene expression between fibroblasts located inside the EMT niche vs those outside. Red: genes with more > 5% expression difference, labeled: red genes with absolute fold change > 1.33. **f)** *VEGFA* expression is specific for fibroblasts in the EMT niche (3D volumetric rendering). Gray: tumor bed (3D surface rendering), black box: EMT niche. **g)** EMT niche macrophage biomarkers. Differential gene expression between macrophages located inside the EMT niche vs those outside. Red: genes with more > 5% expression difference, labeled: red genes with absolute fold change > 1.33. **h)** *SPP1* expression is specific for macrophages in the EMT niche (3D volumetric rendering). Gray: tumor bed (3D surface rendering), black box: EMT niche. **i)** A multicellular EMT niche molecular signature links the EMT niche with reduced survival. Survival plot of 503 lung adenocarcinoma patients from The Cancer Genome Atlas (bulk RNA-seq) stratified into high and low enrichment for the combined expression of *LGALS1*, *NDRG1*, *IGFBP5, VEGA* and *SPP1*.

In light of *LGALS1* role in microenvironment remodeling, we investigated whether the cell type composition of the ‘EMT niche’ differed from the rest of TME. While tumor cells were the most abundant cell type, they composed only 33.1% of the ‘EMT niche’ (**Fig 4d**).

Fibroblasts and macrophages were the main enriched cell types (**Ext data fig4b**) and, together with tumor cells, formed 83.8% of the ‘EMT niche’. We thus focused on fibroblasts and macrophages differentially expressed genes to identify tumor-extrinsic molecular programs active in the ‘EMT niche’.

Fibroblasts upregulated *IGFBP5* and *VEGFA* (**Fig 4e**) and their expression was highly specific to the ‘EMT niche’ (**Fig 4f**, **Ext data fig 4c**). *IGFBP5* (Insulin Growth Factor Binding Protein-5) acts as a reservoir of IGF ligands in the ECM [31] and, especially in the presence of FN1, potentiates IGF signaling, a key pathway promoting cancer cell EMT, proliferation and stemness [32]. Therefore, the local upregulation of *FN1* (Fibronectin-1) and downregulation of *IGFBP7* (Insulin Growth Factor Binding Protein-7), which blocks IGF1R activation [33], could further contribute to the local stimulation of pro-invasive IGF signaling by fibroblasts in the ‘EMT niche’ (**Ext data fig 4d-e**).

At the same time, *VEGFA* (Vascular Endothelial Growth Factor) is a key factor in orchestrating TME remodeling, promoting angiogenesis and the recruitment of tumor-associated macrophages [34]. While the density of capillaries– marked by vascular endothelium and pericytes - was among the lowest in the TME (**Ext data fig4f**), macrophage density peaked in the ‘EMT niche’ (**Ext data fig4g**). Interestingly, macrophages upregulated the expression of *VIM* - expressed by activated macrophages [35]- and the M2 polarization marker *GLUL* (Glutamine synthase) [36] (**Fig 4g**), compatible with a pro-tumorigenic phenotype. In addition, the expression of *SPP1* (Secreted Phosphoprotein-1), a potent chemoattractant for both tumor cells and macrophages that induces the growth of metastatic tumors in a preclinical NSCLC model [37], was highly specific to macrophages in the ‘EMT niche’ (**Fig 4h**).

As tumor invasion typically precedes metastasis to distal organs -the main cause of cancer-related death -, we went on to investigate whether the molecular programs localized to the ‘EMT niche’ were linked with patient survival in a large NSCLC cohort. We leveraged the paired transcriptomic (bulk RNAseq) and survival data for 503 lung adenocarcinoma patients in The Cancer Genome Atlas to investigate the prognostic role of genes specifically expressed in the ‘EMT niche’ (difference of expressing cells > 25%). While single gene enrichments identified patients with a worse prognosis (hazard ratios HR >1), their effects were never statistically significant (**Ext data fig4h-n**). On the other hand, a multicellular signature combining tumor (*LGALS1* and *NDRG1*), fibroblast (*IGFBP5* and *VEGFA*) and macrophage specific (*SPP1*) genes robustly identified patients with shorter overall survival (**Fig 4i**).

Furthermore, *NDRG1* was recently identified as the most specific marker of early brain metastasis (<10 months after diagnosis) in NSCLC patients [38], which matches the clinical history of the patient under study (brain metastasis 11 months after diagnosis).

Taken together, the ‘EMT niche’ represents a specialized compartment enriched with pre-invasive tumor cells, fibroblast and macrophages. Strikingly, the combination of multicellular molecular programs spatially restricted to this niche identified patients with poor prognosis, linking this niche with tumor progression.

### 7. Fibroblast molecular states are linked with the extracellular matrix and cellular composition of their 3D neighborhoods

Tissue stiffening is a hallmark of tumorigenesis, promoting both malignant transformation, EMT and invasion [39]. Thus, we investigated whether tissue biomechanical properties could explain the enrichment of mesenchymal-like tumor cells in the ‘EMT niche’. Quantifying the ECM composition in 3D neighborhoods via second harmonic generation imaging (SHG) (**Fig 5a**, Methods), we confirmed the presence of a homeostatic elastin-rich ECM surrounding lung-resident populations, such as basal and alveolar cells, while fibroblasts and plasma cells lived in a collagen-rich (i.e. desmoplastic) ECM (**Ext data fig 5a**). Compatible with ECM degradation, tumor cell neighborhoods featured the lowest elastin and collagen signals. Taken together, while collagen deposition, which is the main determinant of tissue tension [40], may sustain EMT in tumor cells infiltrating the desmoplastic stroma, it cannot explain EMT induction in the ‘EMT niche’, which featured a collagen-poor ECM similar to the rest of the tumor bed (**Fig 5b**).

**Figure 5.**
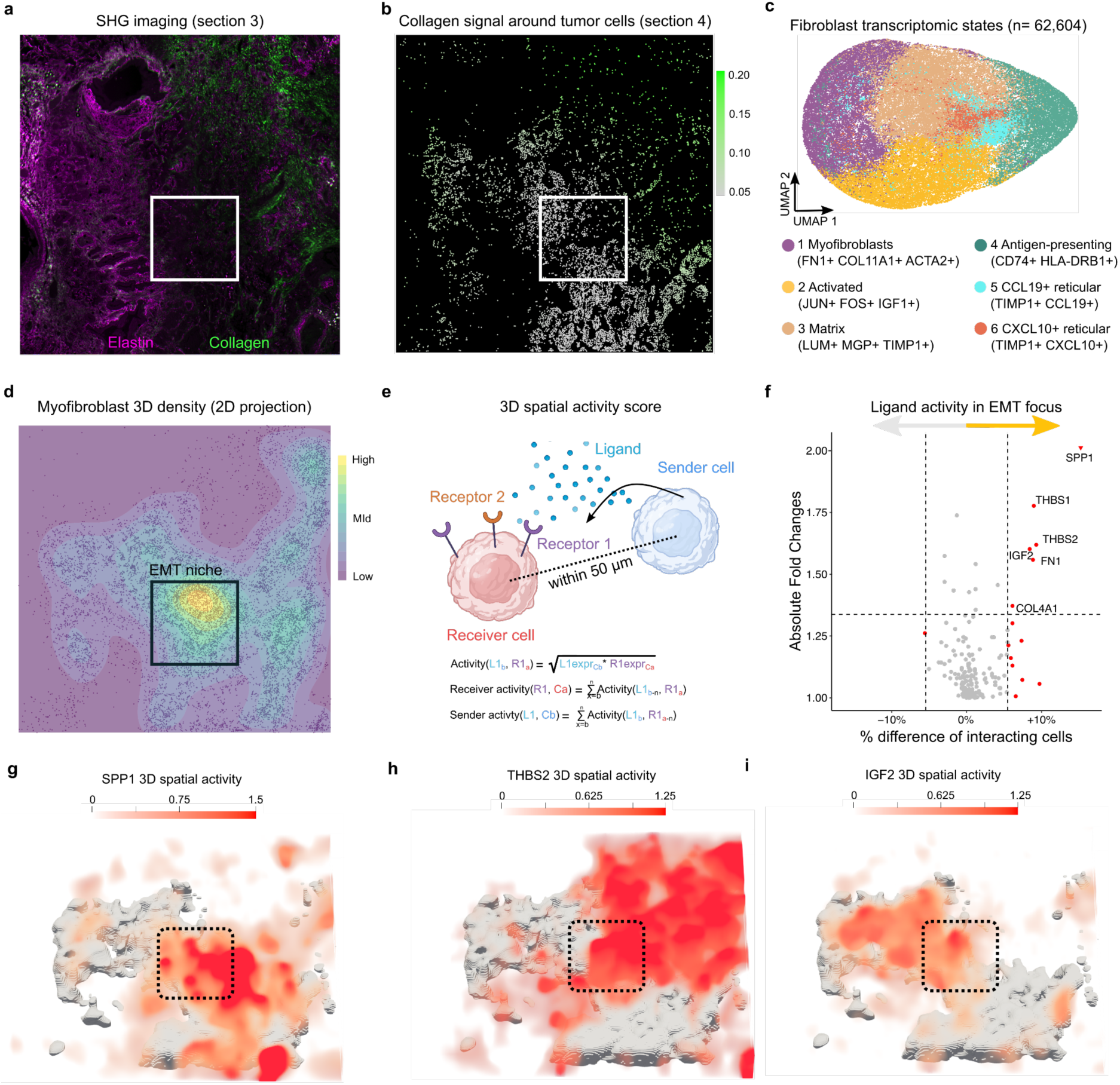
Multimodal analysis of 3D neighborhoods captures the tumor, myofibroblasts and macrophage crosstalk in the EMT niche. **a)** Extracellular matrix (ECM) remodeling in the TME. Second harmonic (SHG) imaging (section 3), purple: elastin, green: fibrillar collagen, white square: EMT niche. **b)** Collagen signal (section 3) in tumor cell neighborhoods (section 4). Tumor cells colored by their mean collagen signal are plotted. white square: EMT niche. **c)** Different fibroblast transcriptomic states in the TME. UMAP plot of fibroblasts colored by cluster assignment. **d)** 3D spatial density plot of myofibroblasts, demonstrating their accumulation in the EMT niche. **e)** 3D spatial analysis of cellular interactions. The sensitive co-detection of ligands and its receptors in 3D cellular neighborhoods is leveraged to quantify the 3D spatial activity scores. **f)** EMT niche-specific ligands. Comparison of 3D spatial activity scores between cells located inside the EMT niche vs those outside. Red: Ligands with more > 5% interaction difference, labeled: red genes with absolute fold change > 1.33. **g-h)** The combination of SPP1 (g), THBS2 (h) and IGF2 (i) signaling is specific to the EMT niche. Black box: EMT niche.

Searching for other factors that could induce/sustain tumor EMT, we shifted our focus towards fibroblasts, which were abundant and upregulated pro-invasive molecular programs in the ‘EMT niche’. The unbiased study of molecular phenotypes for 62,604 fibroblasts revealed six transcriptomic states (**Fig 5c**), which we annotated through literature-informed review of their marker genes (**Ext data fig5b**). Multimodal profiling linked fibroblast molecular states with the mechanical properties and cellular composition of their 3D neighborhoods (**Ext data fig5c-e**). For example, *JUN*+ *FOS*+ *IGF1+* ‘activated fibroblasts’ were enriched in the desmoplastic stroma and surrounded by a collagen-rich ECM, while *CD74*+ *HLA-DRB1*+ ‘’antigen-presenting’, *TIMP1+ CCL19*+ ‘reticular’ and *TIMP1+ CXCL10*+ ‘reticular’ fibroblasts lived in a homeostatic ECM and were spatially enriched in immune niches. On the other hand, *FN1*+ *COL11A1*+ *ACTA2*+ ‘myofibroblasts’ were surrounded by a collagen- and elastin-depleted ECM. Strikingly, they were spatially restricted the tumor bed and their density peaked in the EMT niche (**Fig 5d**). Compared to other fibroblast states, myofibroblasts upregulated ‘EMT niche’ secreted factors (*IGFBP5* and *VEGFA*), ECM-remodeling (*FN1* and *COL11A1*) and smooth muscle-specific contractile proteins (*TAGLN* and *ACTA2)* (**Ext data fig 5b,f**). Similar to wound healing [41], myofibroblast contraction - rather than fibrillar collagen deposition - could mediate the local increase in tissue stiffness. Taken together, myofibroblast density distinguished the ‘EMT niche’ from the rest of the tumor bed, suggesting their central role in tumor invasion.

### 8. 3D cellular neighborhoods capture the tumor, myofibroblasts and macrophage crosstalk in the EMT niche

We therefore shifted our focus to the crosstalk between tumor cells, fibroblasts and macrophages to investigate how the ‘EMT niche’ is formed and maintained. Through the combination of 3D cellular neighborhoods and the sensitive detection of 165 ligands and 144 receptors via ISH (1-2 molecules/cell [14]), we quantified the spatial activity for 480 receptor-ligand pairs in 3D cellular neighborhoods (**Fig 5e,** Methods). In the CellChat database [42], interactions enriched in the ‘EMT niche’ (**Fig 5f**) belonged either to the ‘ECM-Receptor’ category, including FN1 (Fibronectin-1), THBS1 (Thrombospondin-1), THBS2 (Thrombospondin-2) and COL4A1 (Collagen type IV, alpha-1 chain), or were annotated as ‘Secreted signaling’, including SPP1 (Osteopontin) and IGF2 (Insulin-like growth factor-2).

Studying the directionality of these signaling networks in 3D neighborhoods, we confirmed the role of macrophages as source of SPP1 signaling and identified fibroblasts and tumor cells as main senders of ‘ECM-receptor’ and IGF2 interactions, respectively (**Ext data fig5g**, orange dots). Strickingly, ‘ECM-receptor’ interactions were mainly received by tumor cells and converged on multiple integrin receptors, including ITGA3, ITGAV and ITGB1 (**Ext data fig5g**, blue dots). Integrins are the main cellular adhesion receptors and their central role in tumor invasion is well-established. They represent key components of the cell migration machinery and their engagement activates intracellular pathways promoting survival, EMT and invasion [43]. Accordingly, although individual signaling could be also found in other niches, the combination of SPP1, FN1, IGF2 and THBS2 spatial activity was collectively restricted to the ‘EMT niche’ and distinguished it from the rest of the tumor bed (**Fig 5g-i, Ext data fig 5h-l**). Interestingly, IGF signaling promoted myofibroblast differentiation of lung fibroblasts grown on a soft substrate [44], [45]. Intriguingly, tumor-derived IGF2 may promote myofibroblast accumulation in the EMT niche, which further potentiates pro-invasive IGF signaling through IGFBPs and FN1 production. Furthermore, the co-localization of THBS2+ myofibroblasts and SPP1+ macrophages was recently linked with T-cell exclusion in both lung [42] and colorectal tumors [46], linking the EMT niche with the remodeling of the tumor immune microenvironment.

### 9. 3D neighborhoods boost the quantification of immune niche-specific interactions, including druggable immune checkpoints

Under the selective pressure of the immune system, tumor cells eventually develop strategies to suppress anti-tumoral immune responses, including the reprogramming of antigen-presenting cells (APCs, i.e. macrophages and dendritic cells). In turn, APCs may promote the recruitment of regulatory T cells, further restricting the activation of tumor-specific cytotoxic T cells [47]. Compatible with tumor immune escape, APC and regulatory T cells were co-enriched in ‘macrophage niches’ and ‘dendritic cell niches’ (**Fig 6a**), while cytotoxic T cells - despite their abundance-did not infiltrate the tumor bed and accumulated in ‘T cell niches’ (**Fig 6a,b**).

**Figure 6.**
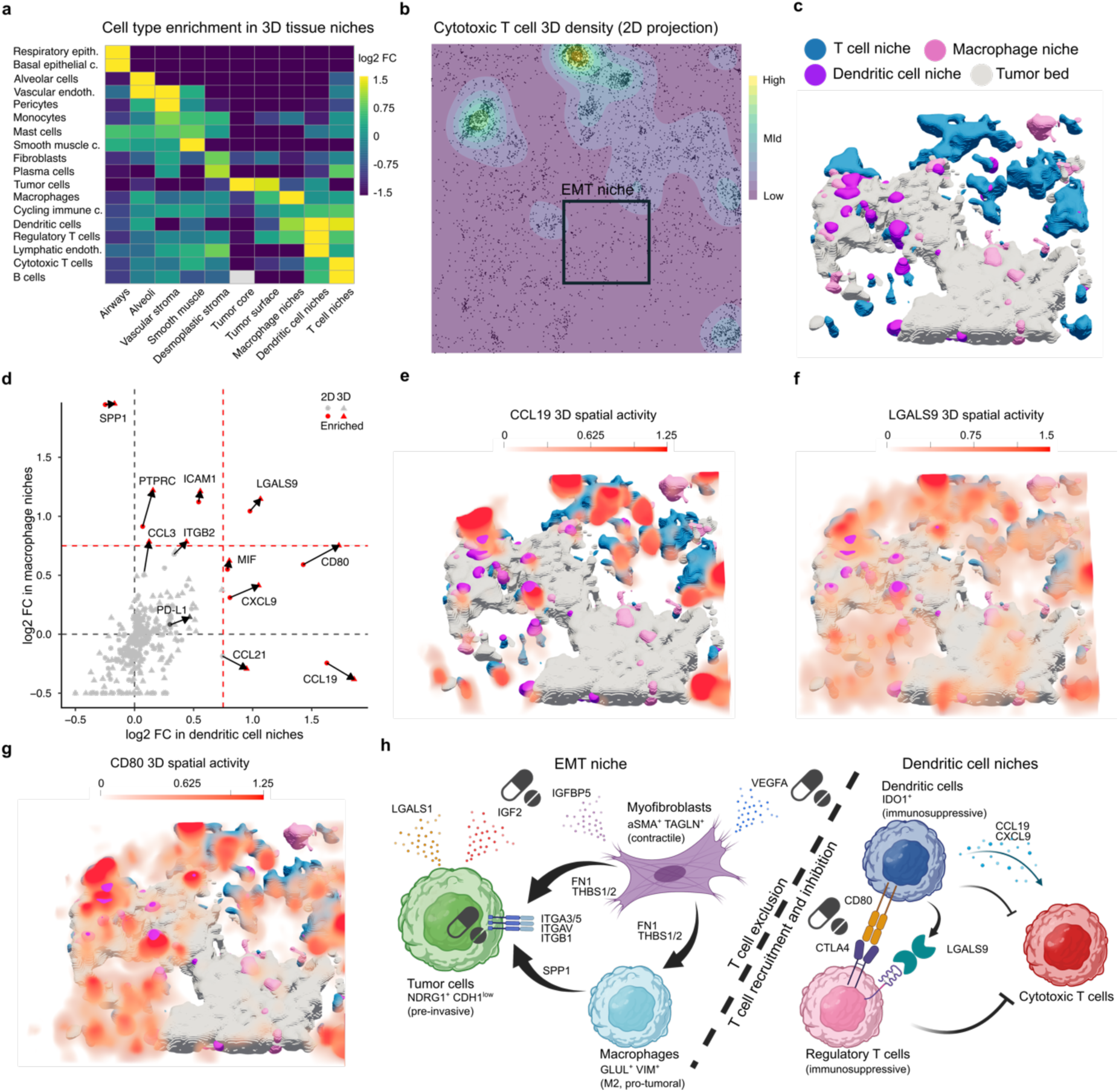
3D neighborhoods boost the identification of immune niche-specific interactions, including druggable immune checkpoints. **a)** Heatmap showing enrichment of cell types in 3D niches. log2 fold changes are computed from the ratio between the observed and expected number of cells per cluster in each 3D niche. **b)** 3D spatial density plot of cytotoxic T cells, demonstrating their accumulation in immune niches and exclusion from the tumor bed. **c)** Spatial organization of immune niches in the TME. 3D rendering of ‘Dendritic cell niches’ (purple), ‘Macrophage niches’ (magenta), ‘T cell niches’ (blue) and the tumor bed (gray). **d)** Systematic scoring of interactions identifies ligands enriched in immune niches. log2 Fold changes (log2 FC) for 164 ligands in ‘Dendritic cell niches’ (x axis) and ‘Macrophage niches’ (y axis). FC are computed comparing average ligand sender scores for cells inside immune niches against those outside. 11 ligands with a log2FC > 0.75 in either niche (red) and PD-L1 (dark grey) are labeled. Arrows connect 2D (dots) and 3D (triangles) enrichment scores for each enriched ligand and PD-L1. **e-g)** CCL19 (e), LGALS9 (f) and CD80 (g) signaling is enriched in immune niches. **h)** Cellular interactions driving tumor invasion and immune escape, revealing mechanism-based personalized targets for cancer interception. Summary scheme of selected molecular features and interaction in the ‘EMT niche’ (left) and ‘Dendritic cell niche’ (right). The pill symbol indicates druggable interactions.

To identify immunomodulatory interactions, we focused on interactions that distinguished ‘dendritic cell niches’ and ‘macrophage niches’ from the rest of the TME (**Fig 6c**).

Compatible with their role in immune cell recruitment (i.e. chemotaxis), numerous chemokines were enriched in immune niches. The 3D spatial activity CCL19 (CC-chemokine ligand 19), CCL21 and CXCL9 (C-X-C motif ligand 9), which regulate the homing of dendritic and T cells in lymphoid tissues [48], marked location of ‘dendritic cell niches’ and ‘T cell niches’ in the TME (**Fig 6e**, **Ext data fig 6b-c, Supp Video 2**). In line with their chemotactic role, CCL19 and CXCL9 interactions were received by most cell types in ‘dendritic niches’, with the exception of tumor cells (**Ext data fig 6d**), suggesting their involvement in the induction of - rather than their recruitment to - ‘dendritic niches’. Tumor cells indeed acted as main MIF (Macrophage Inhibitory Factor) senders, a potent chemoattractant upregulated in several tumors, including NSCLC [49]. MIF, whose spatial activity was widespread in the tumor bed (**Ext data fig 6e**), promotes immunosuppressive myeloid reprogramming and its inhibition decreased regulatory T cell and promoted cytotoxic T cell infiltration in a melanoma lung metastasis model [50].

At the same time, LGALS9 (Galectin 9) and CD80 emerged as the ligands with the strongest shared upregulation across all immune niches (**Fig 6f-g**), highlighting their importance in regulating anti-tumoral immune responses. While the network of LGALS9 interactions in ‘dendritic cell niches’ involved all cell types, CD80 interactions mainly occurred between dendritic cells and CTLA-4+ (Cytotoxic T-Lymphocyte Antigen 4) regulatory T cells. While LGALS9-CD44 interactions promote the persistence and function of regulatory T cells in the TME [51], CTLA4-CD80 interactions mediate their immunosuppressive function [52]. In fact, CTLA4 prevents the interaction of CD80 with the co-stimulatory receptor CD28 on cytotoxic T cell [53] and promotes the expression of IDO (indoleamine-2,3-dioxygenase) by dendritic cells, which deprives adjacent T cells of tryptophan and further inhibits their activation [54]. Confirming the functional activity of the CD80-CTLA4 axis, dendritic cells engaged in CD80-CTLA4 interactions upregulated the expression of IDO1 (**Ext data fig6f**).

Remarkably, 3D neighborhoods consistently returned higher fold changes compared to their 2D counterparts, thereby boosting the identification of enriched ligands across all immune niches (**Fig 6d**, **Ext data fig 6a**).

Taken together, 3D neighborhoods identified signaling networks spatially linked with tumor invasion and immune responses and may provide the molecular rationale for their therapeutic blockade, especially in the era of personalized medicine and mechanism-based combination therapies [55]. Importantly, monoclonal antibodies targeting key receptors, such as anti-CTLA4 or anti-VEGF agents, are already approved for the treatment of NSCLC, while numerous inhibitors blocking IGF receptors, integrin signaling, or IDO enzymatic activity are currently being evaluated in clinical trials (**Fig 6h**).

## DISCUSSION

This study represents the proof-of-principle for the molecular reconstruction of 3D cellular neighborhoods combining high-plex, single-cell resolved spatial transcriptomic data with imaging readout of tissue biomechanical properties. 3D neighborhoods captured the tissue organization of cell types in multicellular niches and enabled the exploration of niche-specific molecular states. Furthermore, the spatial mapping of more than 450 receptor-ligand pairs in 3D cellular neighborhoods allowed the systematic and highly sensitive (1-2 molecules/cell [14]) quantification of niche-specific multicellular crosstalk. Leveraging the close spatial proximity of interacting cells in 3D, we focused on physically possible interactions, thereby boosting the specificity of our approach. Finally, the alignment of sections profiled with multiple modalities allowed the integrative analysis of tissue gene expression, ECM composition and protein marker levels. For example, combining SHG with cellular communication analysis was instrumental in interpreting how myofibroblasts – together with macrophages-modulate pro-invasive integrin signaling on tumor cells in the EMT niche.

Furthermore, the integration of mechanical and molecular readouts revealed how fibroblast states in the TME are linked with the cellular and ECM composition of their neighborhoods. In general, given the profound impact of tissue biomechanics in shaping cellular molecular states (and *vice versa*), we envision that the systematic integration of spatial mechanical and molecular readouts will reveal novel insights into tissue functioning in health and disease (e.g., shedding light on the intricate role of tumor-stromal interplay).

While bi-dimensional histology and spatial omics [56], [57] represent powerful methods for the description of tissue pathology and the discovery of physiological and pathological cellular interplay, our data argue that 3D neighborhoods improved the identification and characterization of tissue niches. In line with a recent colorectal cancer study [7], 3D neighborhoods captured the spatial continuity of seemingly disconnected ‘T cell niches’ in 2D. Furthermore, the study of ‘dendritic cell niches’, where local anti-tumoral immune responses are orchestrated at the tumor surface was only possible in 3D. Importantly, 3D neighborhoods boosted the quantification of cell-cell interactions, which is central to coordination of immune responses in health and disease states [58]. Here we defined 3D neighborhoods using a 50 µm radius and a 30µm section-to-section gap. Future studies will be required to identify both the most cost-effective gap maximizing 3D information while minimizing repeated sampling of the same cells (average cell radius 10-20 micron) and which neighborhood radius is most appropriate to capture short vs long-distance molecular dependencies. Nevertheless, taken together, our data demonstrate the importance of 3D neighborhoods for the study of immune niches in the TME.

Focusing on an early-stage, aggressive NSCLC tumor, we demonstrate the ability of 3D neighborhoods to capture the molecular mechanisms linked with tumor invasion and immune escape. While a recent IMC study demonstrated how the TME spatial organization can accurately predict survival in early-stage NSCLC patients [59], limited plexity prevented the identification of personalized drug targets. In this study, 3D neighborhoods spatially linked VEGF, IGF and integrin signaling with tumor invasion in the ‘EMT niche’ and highlighted CTLA4 and IDO1 as key immunosuppressive molecules in the patient under study. These could serve as biomarkers for the response to agents already approved for NSCLC therapy, including anti-VEGF (e.g. bevacizumab) and anti-CTLA4 agents (e.g. ipilimumab) monoclonal antibodies. Personalized biomarkers not only have a crucial role in increasing therapeutic efficacy and limiting treatment-related costs and toxicities but are also urgently needed for the clinical study of novel therapeutic approaches. For instance, the failure of anti-IGF agents - after decades of encouraging preclinical results [53] – is mainly attributed to recruitment of unselected NSCLC cohorts in phase III clinical trials due to the lack of predictive biomarkers [60]. Like the simultaneous identification of multiple druggable mutations by NGS sequencing opened the door to combination therapies [19], we hypothesize that the high-plex quantification of active molecular pathways will pave the way to effective, personalized combination therapies. In this regard, small molecule inhibitors blocking the signal transduction pathways downstream of multiple receptors are particularly promising. For example, the simultaneous inhibition of integrin signalling (via FAK inhibitors) and VEGFR [61], [62] or IGFR receptors [62] could represent a particularly attractive therapeutic strategy in the patient under study. Ultimately, interventional studies enrolling large patient cohorts will be critical to assess the predictive ability and clinical benefit of the mechanism-based, personalized biomarkers we identified in this single-patient, observational study.

Furthermore, while here we demonstrate the ability of a large cancer-related panel to profile ligand-receptor interactions in well-established signaling pathways, genome-wide assays will ultimately be needed to unbiasedly profile cell-cell interactions and discover novel mechanisms and therapeutic targets. At the same time, unbiased assays will enable the simultaneous profiling of ligand, receptors and intracellular response genes, further increasing the likelihood of identifying active signaling axes. While mRNA represents a sensitive, antibody-independent readout for protein production, discrepancies between transcripts and protein levels do exist. Therefore, the systematic understanding of the molecular architecture of the TME will likely require the simultaneous profiling of proteins and transcripts in single cells. Finally, the integration of additional data modalities, such as epigenomics or metabolomics, will provide an even more complete picture.

## Supporting information

Video abstract

## Acknowledgements

We thank Enes Senel, Marvin Jens, Leon Strenger, Kamil Lisek, Lieke van de Haar, Marie Schott, Cledi Alicia Cerda Jara, all other members of the Rajewsky lab for critical and useful discussions. Alexandra Tschernycheff, Veronica Jakobi and Margareta Herzog for the lab organizational support. We thank Anca Margineanu of the Advanced Light Microscopy Technology Platform at the Max-Delbrück-Center for her technical support with two-photon microscopy. A special thanks to Gaetano Gargiulo for reviewing the manuscript. N.R. and T.M.P. thank Nir Friedman for discussions and Stan Gorski for editorial feedback. Illustrations in Figures 1a and 6h and Extended data figure 1a were created with Biorender.

## Author Contributions

Conceptualization and design - T.M.P., I.L., N.K. and N.R. Methodology - T.M.P., N.K. Sample collection and pathology analysis - S. S. Deep learning HE image analysis - S.S., L.R., G.D. Sample processing and quality control - G. T., A. B. Spatial transcriptomics data collection and preprocessing - Y. K., S. K., S. M., M. G. Second harmonic imaging data collection and preprocessing - A.W. Immunofluorescence data collection and preprocessing - D. L.-P., S. F., J. N., P.J. Computational analysis - T.M.P., N.K. 3D renderings: D. L.-P. Writing – Original Draft: T.M.P. Writing – Review & Editing: T.M.P. and N.R. with input from I.T., I.L., M.C., S.P., F.C., N. K. and F., K. All Authors discussed and analyzed the data.

## MATERIALS AND METHODS

### Study design

To study three-dimensional (3D) cellular neighborhoods in the tumor microenvironment (TME), we focused on a formalin-fixed, paraffin-embedded (FFPE) tumor block from a NSCLC patient and collected 34 5 µm-thick consecutive sections using a microtome. We processed sections 4, 10, 16, 22, 28 and 34 with CosMx (section-to-section distance 30 µm) to generate high-plex, single-cell resolved spatial transcriptomics data, section 3 with both second harmonic imaging (SHG) to study extracellular matrix (ECM) composition and hematoxylin and eosin (H&E) staining to capture tissue morphology and section 12 with immunofluorescence (IF) to validate tumor epithelial-to-mesenchymal transition (EMT) at the protein level. We then employed computational methods for the 3D alignment of the analyzed sections to generate a 3D multimodal atlas of NSCLC at single cell resolution.

### NSCLC patient’s characteristics

The patient was a 63-year-old female, who presented with a pulmonary tumor mass in the apex of the right upper lobe (RUL) in the positron emission tomography in March 2020. Regarding smoking status, she was a 40 pack-year ex-smoker. The patient was fully active, without any physical restrictions (ECOG grade 0). A transbronchial lung biopsy revealed a TTF1-positive lung adenocarcinoma (LUAD) with acinar morphology. Staging workup by abdominal and a brain CT-scan showed no other potential lesions, additionally the bone scintigraphy was negative. Two months after initial diagnosis, a resection of the RUL was performed. The pathological examination revealed a tumor with a maximum diameter of 23 mm, infiltration of the visceral pleura, lymphovascular invasion, and three metastases into hilar lymph nodes with a maximum diameter of 7 mm. The tumor was completely resected. The pathological tumor classification was as follows: pT2a pN1 (3/24) L1 V0 Pn0 G2 R0. The patient received three cycles of an adjuvant-combined chemotherapy (Cisplatin + Vinorelbine).

In February 2021, the patient started having neuronal symptoms and a magnetic resonance tomography of the head was performed revealing a 10 mm tumor in the frontal cortex. The tumor was resected and a metastasis of the LUAD was histologically confirmed. Afterwards the patient received cranial radiotherapy in the form of a volumetric intensity modulated arc therapy (VMAT) at a dosage of 25,6 Gray (Gy). In August 2021 a second metastasis was diagnosed in the left upper lobe (LUL) of the lung.

### Sample collection and histological examination

Resected specimens and core needle biopsies were fixed in 10% buffered formalin before gross processing. After overnight fixation, the specimens were cut in 5-mm-thick slices. As a first step, the tumor was detected, described and the tumor diameter as well as the minimum distances to the visceral pleura and the resection margins of lung parenchyma and bronchus were measured. Next, the resection margins and the representative tumor parts showing the relation to the relevant anatomical structures, described above, were embedded. Furthermore, we retrieved all macroscopic detectable lymph nodes. Subsequently the tissue or biopsies were embedded in paraffin and were stored at room temperature at the archive of the Institute of Pathology at the Charité University Hospital, Campus Mitte. Histological examination, including diagnosis, tumor grading, pTNM-classification, angioinvasion, lymphatic invasion, and tumor stage was done according to the 8th edition of the TNM classification (AJCC). The study was performed according to the ethical principles for medical research of the Declaration of Helsinki and approval was approved by the Ethics Committee of the Charité University Medical Department in Berlin (EA4/243/21).

### Deep learning segmentation of whole slide H&E images

For learning a tissue segmentation model, we collected around ∼5,000 representative pathologist annotations for 4 morphological sub-categories on H&E tissue morphology: “Carcinoma”, “Stroma”, “Necrosis”, and “Normal lung” (e.g. including “Alveoli”, “Capillaries”, “Respiratory epithelium”, “Vessels”, etc.) for model training. For the segmentation model, we used a U-Net [63] architecture with a ResNet101 backbone [64]. We trained models over various hyperparameters for 50 epochs using the Adam optimizer [65] and selected the top five models based on the global F1 performance on a validation set. We then combined these five models into a mean ensemble, which achieved a global F1 performance of ∼0.93 on a hold-out test set.

### Quantification of RNA fragmentation

To quantify the extent of RNA fragmentation, we collected 3 10 µm sections using a Microtome (Leica Byostems). We then extracted RNA using Qiagen RNAse FFPE kit according to manufacturer instructions and evaluated the percentage of total RNA fragments longer than 200 nucleotides (DV200) using the TapeStation (Agilent). Compatible with formalin fixation and prolonged storage at room temperature, the DV200 score was 60%.

### CosMx sample processing, staining, imaging, and cell segmentation for 1000-plex RNA profiling

CosMx sample processing, staining, imaging, and cell segmentation were performed as previously described [14]. Briefly, tissue sections were placed to VWR Superfrost Plus Micro Slide (Cat# 48311-703) for optimal adherence. Slides were then dried at 37°C overnight, followed by deparaffinization, antigen retrieval and proteinase mediated permeabilization (https://nanostring.com/products/cosmx-spatial-molecular-imager/single-cell-imaging-overview/). 1 nM RNA-ISH probes were applied for hybridization at 37°C overnight. After stringent wash, a flow cell was assembled on top of the slide and cyclic RNA readout on CosMx was performed (16-digit encoding strategy). After all cycles were completed, additional visualization markers for morphology and cell segmentation were added including pan-cytokeratin, CD45, CD3, CD298/B2M, and DAPI. Twenty-four 0.985mm × 0.657mm fields of view (FOVs) were selected for data collection in each slice. The CosMx optical system has an epifluorescent configuration based on a customized water objective (13×, NA 0.82), and uses widefield illumination, with a mix of lasers and light-emitting diodes (385 nm, 488 nm, 530 nm, 590 nm, 647 nm) that allow imaging of DAPI, Alexa Fluor-488, Atto-532, Dyomics Dy-605 and Alexa Fluor-647, as well as removal of photocleavable dye components. The camera was a FLIR BFS-U3_200S6M-C based on the IMX183 Sony industrial CMOS sensor (pixel size 180nm). A 3D multichannel image stack (9 frames) was obtained at each FOV location, with the step size of 0.8 μm. Registration, feature extraction, localization, decoding of the presence of individual transcripts, and deep-learning based cell segmentation (developed upon Cellpose [66]) were performed as previously described [14]. The final segmentation mapped each transcript in the registered images to the corresponding cell, as well as to subcellular compartments (nuclei, cytoplasm, membrane), where the transcript is located.

### Transcriptomic clustering of segmented cells

For downstream analyses we used the package Seurat [67] (v4.0.4) in R (v4.1, https://www.R-project.org/). For each section, we imported 3 matrices containing the gene expression, metadata and positions of segmented cells. We removed the negative probes from the gene expression matrix, defined a unique cell name and created a merged Seurat object with data from all the sections.

To identify cell types present in the tumor microenvironment, we adopted a very conservative filtering strategy removing only cells with less than 10 detected genes and removing genes detected in less than 1 cell. We then computed SCT-normalized and scaled gene expression counts [68] and computed the 50 most variable principal components (PCs). We selected the first 30 PCs to create a shared nearest neighbor graph and to compute a two-dimensional UMAP plot [69] used for data visualization in a low dimensional space. Finally, we partitioned the shared nearest neighbor graph using a resolution of 0.8 and identified 24 transcriptomic clusters.

### Cell type annotation of transcriptomic clusters

To annotate cell types present in the tumor microenvironment, we identified marker genes enriched in each cluster for knowledge-based cell type annotation. Epithelial clusters, characterized by positivity to panCK immunofluorescent staining, comprised both tumor cells (*EPCAM+ S100A10*+) and non-malignant epithelial cells, including respiratory epithelium (*SCGB3A1*+), basal epithelial cells (*KRT5*+) and alveolar cells (*FGG*+). We also detected numerous stromal populations, including fibroblasts (*COL1A2*+), vascular (*PECAM1*+) and lymphatic (*FABP5*+) endothelial cells, pericytes (*PDGFRB*+) and smooth muscle cells (*TAGLN*+). We identified innate and adaptive immune cell populations, including macrophages (*CD68*+), monocytes (*LYZ*+), dendritic cells (*LAMP3*+), mast cells (*KIT*+), cytotoxic (*CD8A*+) and regulatory (*FOXP3*+) T cells, B cells (*IGHM*+), plasma cells (*CD79A*+) and cycling immune cells (*MKI67*+). Finally, we merged clusters with similar marker gene expression as probably related to different cellular states of the same cell type. We thus annotated clusters ‘0’ and ‘1’ as fibroblasts, clusters ‘2’ and ‘7’ as macrophages and clusters ‘4’, ‘5’, ‘20’, ‘22’ and ‘23’ as tumor cells.

### Data integration with reference atlases

To compare gene expression profiles with published reference atlases, we performed label transfer using the standard Seurat pipeline. We integrated our data with the Human Lung Cell Atlas [21] to annotate lung resident cell types and with a NSCLC single cell cohort [22] to annotate tumor-specific cell types. For each reference dataset, we identified a subset of 900 shared ‘features’ (i.e. genes) using the SelectIntegrationFeatures function (nfeatures = 900) and pairs of ‘anchors’ (i.e. cells) between the reference and our query dataset using the FindTransferAnchors function (normalization.method = “SCT”, reference.assay = ‘SCT’, query.assay = ‘SCT’, reduction = ‘pcaproject’, dims = 1:20, features = features, nn.method= ‘rann’, eps = 0.5). Then, we leveraged the TransferData function (anchorset = anchors, prediction.assay = TRUE, weight.reduction = ‘pcaproject’, dims = 1:20, eps = 0.5) to score each cell in our query dataset for similarity with annotated cell types in the reference dataset.

### 3D alignment of spatial transcriptomic data

To align high-plex, single-cell resolved spatial transcriptomic data from 6 sequential, non-consecutive sections, we leveraged the Spatial Transcriptomics ImgLib2/Imaging Project (STIM) [23]. With STIM, we first converted our spatial transcriptomics data to the n5 image format for efficient storage and processing using the ‘st-resave’ function. In doing so, we assigned a channel to each gene and modeled gene expression values as pixel intensities at the center of the segmentation mask. Then, we applied the ‘st-align-pairs’ function to align each section to the one above and below (r=1) using the Scale Invariant Feature Transform (SIFT) [70] according to the expression of the 15 genes with highest standard deviation (n=15). Finally, we applied the ‘st-align-global’ function to identify a global optimum that minimizes the distances between all corresponding points across all pairs of slices (--absoluteThreshold 100 --sf 0.5 --lambda 0.5 --skipICP).

### Identification of 2D and 3D cellular neighborhoods

Cellular neighborhoods in 2D and 3D were computed with a custom Python script as follows. First, for a given cell, the Euclidean distances in 2 or 3D between that cell and all other cells in the dataset were computed. This set of distances was then filtered to remove distances greater than r= 50 μm, resulting in a list of neighboring cells for that given cell. This list was finally used to construct the 2D and 3D neighborhood matrices by counting the number of cells for each of the 18 cell types present in each cellular neighborhood.

### 3D visualization of airways and 3D cellular neighborhoods

To generate 3D plots at single cell-resolution, we leveraged the Giotto R package (v1.1.2) [71] and the Plotly R package (v4.10.1, https://plotly-r.com). Briefly, we first created a Giotto object with the 3D cell centers coordinates and cell type annotations. To visualize airways, we used the ‘spatPlot3D’ function (a wrapper for Plotly 3D scatter plot) highlighting ‘respiratory epithelium’ cells in green and showing all other cell types in gray. To visualize 3D cellular neighborhoods, we selected an exemplary center cell (Section 22, FOV 1, Segmentation ID 132) and plotted neighboring cells within a 100 µm center-to-center distance. We then generated a 3D scatter plot using Plotly, highlighting the center cell in red, coloring the cells within a 50 µm according to their cell type and showing the cells outside of the 3D neighborhood (between 50 and 100 µm) in gray.

### Identification and annotation of 2D and 3D multicellular niches

To identify 2D/3D multicellular niches in the tumor microenvironment, we imported the 2D/3D neighborhood matrix as a new assay in Seurat. We excluded cells with incomplete 3D neighborhoods, namely those located in sections 4 and 34 and those within 50µm from the edges of sections 6, 12, 18 and 24. We then performed UMAP dimensionality reduction and clustering (resolution = 0.3). 3D neighborhoods analysis returned 13 clusters (**Supp Fig 1a**). Furthermore, we merged clusters ‘3’, ‘10’ and ‘12’ as ‘T cell niches’ and clusters ‘5’ and ‘9’ as ‘airways’ given their shared neighborhood composition and spatial patterns (**Supp Fig 1b-c**). In this way, we identified a total of 10 unique 3D niches, which we annotated based on the average cell type counts per 3D neighborhood: ‘tumor core’ (Tumor cells^high^, Fibroblasts^low^, Macrophages^low^), ‘tumor surface’ (Tumor cells^high^, Fibroblasts^mid^, Macrophages^mid^, Cytotoxic T cells^low^), ‘airways’ (Respiratory epithelium^high^, Basal epithelial cells^high^), ‘alveoli’ (Alveolar cells^high^), ‘desmoplastic stroma’ (Fibroblasts^high^, Plasma cells^low^), ‘vascular stroma’ (Fibroblasts^mid^, Vascular endothelium^mid^, Pericytes^mid^), ‘smooth muscle’ (Smooth muscle cells^high^), ‘Macrophage niche’ (Macrophages^high^, Tumor cells^mid^), ‘Dendritic cell niche’ (Dendritic cells^high^, Tumor cells^mid^, Macrophages^mid^, Cytotoxic T cells^mid^, Regulatory T cells^low^) and ‘T cell niche’ (Cytotoxic T cells^high^). At the same time, 2D neighborhoods analysis returned 15 clusters (**Supp Fig 1d**). Similarly, we merged clusters ‘8’, ‘10’ and ‘13’ as ‘tumor core’, clusters ‘2’ and ‘3’ as ‘tumor surface’, clusters ‘5’ and ‘12’ as ‘airways’ and clusters ‘4’ and ‘14’ as ‘T cell niches’ and identified 10 multicellular niches (**Supp Fig 1e**), which overlapped with 3D ones, with the exception of ‘dendritic cell niches’ identified only in 3D and ‘macrophage-rich stroma’ (Fibroblasts^mid^, Macrophages^mid^) identified only in 2D.

### Tumor pseudotime analysis

To reconstruct their molecular dynamics, we ordered tumor cells according to their pseudotime. We first generated a Seurat object including only tumor cells using the subset function and then re-normalized gene expression counts using SCTransform. We then selected the top 400 variable genes and grouped them in 5 PCs using the RunPCA function for downstream analyses. These included UMAP embedding using the RunUMAP function and pseudotime analysis. For the latter, we converted the Seurat object into SingleCellExperiment format using the R package SingleCellExperiment (v1.16.0) [72] and then used the slingshot function of the R package slingshot (v2.2.1) [73] to compute tumor cell pseudotime. Finally, we converted pseudotime scores to ranks (between 0 and 1) and added them as metadata in the Seurat object for plotting.

### Immunofluorescence (IF) imaging

The IF workflow for FFPE tissues was performed as previously described [74]. In detail, FFPE tissue was deparaffinized, rehydrated and a two-step antigen retrieval was performed at pH 6 and pH 9 sequentially with a rinsing step in PBS in between. Slides were blocked in Odyssey blocking buffer (LI-COR BioScience, Cat # 927-70001) for 30 minutes. Prior to antibody incubation, a pre-bleaching step was performed in 4.5% H_2_O_2_ and 24 mM NaOH diluted in PBS for 30 minutes at room temperature in the presence of white light. The tissue was counterstained with Hoechst 33342 (Thermo Fisher Scientific, Cat #62249), mounted with ProLong™ Diamond (Thermo Fisher Scientific, Cat #P36961) and imaged. After acquisition, slides were soaked in PBS for 5-10 minutes to remove the coverslip and antibody incubation was performed in a humid chamber at 4 °C overnight. Slides were then washed, mounted, imaged and bleached. Images were acquired on a Zeiss Axioscan7 Slide scanner using an EC Plan-Neofluor 20x/0.5 M27 objective with 1×1 binning.

### Antibodies

| Antibody target protein | Vendor | Catalog no. | Clone | Fluorophore | Research resource identifier |
| --- | --- | --- | --- | --- | --- |
| E-cadherin | CST | 3199 | 24E10 | Alexa Fluor® 488 | AB_10691457 |
| Pan-Cytokeratin | Thermo Fisher Scientific | 41-9003-80 | AE1/AE3 | eFluor™ 570 | AB_11218704 |

### Preprocessing of imaging data

To stitch and register the .czi image files, acquired on the Zeiss Axioscan7 widefield microscope, the ASHLAR algorithm (v 1.17.0) [75] was used. To correct the tissue autofluorescent background signal, we performed background subtraction. Images were acquired before the first round of antibody staining; using the same settings for each channel (only exposure time was modified). The software package from github.com/SchapiroLabor/Background_subtraction was used (v 0.3.3) based on the following formula:

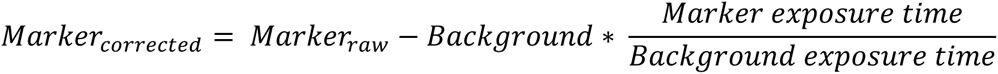

### Registration of IF and CosMx images

The IF multi-channel image stack from Section 12 was registered to the CosMx images of Section 10. To do this, the IF data stored as pyramidal OME-TIFF images were cropped into a rectangular region of interest (ROI) using bftools from Bio-Formats (v6.11.1) [76]. This was done to exclude parts of the tissue outside CosMx ROI, reducing the overall size of the images and simplifying the registration process. The 24 contiguous fields of view of CosMx raw imaging data for Section 10 were then stitched into a single image using the Grid/Collection stitching v1.2 plugin in Fiji v1.53t [77]. The average z-stack projection of the stitched stack was exported as a pyramidal OME-TIFF image. Rigid and affine registration was performed using wsireg 0.3.7 with default parameters, with the full IF stack of Section 12 as the moving image, and the stitched CosMx stack of Section 10 as the fixed (target) image.

### Survival analysis

We downloaded the expression matrices and the associated clinical metadata of the LUAD study directly from the TCGA cohort (https://www.cancer.gov/about-nci/organization/ccg/research/structural-genomics/tcga). We subsetted the cohort to remove patients that received prior treatment, as well as donors for whom only data from healthy tissue was available (control group). The corresponding bulk RNA expression data for these patients was then obtained, and the FPKM values were log-transformed for further analyses. To perform censored survival analysis, we considered the “days to last follow up” to be equivalent to the “days to death”, for the cases where the donor was still alive. To compute gene enrichment scores, we used the R package GSVA v.1.46.0 with the method “zscore” and a threshold of 0 to cluster the donors into low/high expression scores. The survival curve was computed with the functions “Surv” and “survfit” of the R package survival v.3.4.0, and by considering the “days to last follow up” or “days to death” as time, the vital status (alive vs dead) as the event, and stratifying the data by the clusters identified by the gene enrichment analysis as described above. To compute the hazard ratio, we computed a Cox model using the function coxph of the R package survival and inserted the result into the function tbl_regression of the R package gtsummary v.1.7.0.

### Second Harmonic Imaging

Label free imaging of collagen and elastin was performed on a Zeiss LSM 880 NLO equipped with a Plan-Apochromat 10x NA 0.45 objective (Carl Zeiss Microscopy GmbH, Jena, Germany) and a tunable femtosecond titanium-sapphire laser (Chameleon-Ultra II, Coherent, Santa Clara, California). Using an excitation wavelength of 800 nm, the second-harmonic generation signal from collagen was collected through a 395 – 405 nm spectral window on to a GaAsP detector and autofluorescence emission from elastin was collected through a 435 - 480 nm spectral window on to a PMT.

### ECM composition analysis

To quantify ECM levels of collagen and elastin in section 4 cellular neighborhoods, we first imported cropped and aligned section 3 SHG TIFF images in R using the grDevices package (v. 4.1.3). We then normalized the collagen and elastin channels by dividing the pixel intensities by the maximum value in the respective channel. Finally, we quantified the mean collagen and elastin pixel intensities in the square with side 2x 50 µm centered on the segmentation center of each cell in section 4.

### Fibroblast transcriptomic clustering and annotation

To identify transcriptomic states of the fibroblast in the TME, we selected cells annotated as fibroblasts with more than 100 detected transcripts for unsupervised clustering as described above, this time selecting the first 5 PCs and identifying 6 clusters and a resolution= 0.15. We then identified marker genes enriched in each cluster for literature-based cluster annotation [78]–[82]: ‘matrix fibroblasts’ (*LUM*+ *MGP*+ *TIMP1*+), ‘myofibroblasts’ (*FN1*+ *COL11A1*+ *ACTA2*+), ‘activated fibroblasts’ (*JUN*+ *FOS*+ *IGF1+*), ‘antigen-presenting’ (*CD74*+ *HLA-DRB1*+), *‘CCL19*+ reticular’ and *‘CXCL10*+ reticular’ fibroblasts.

### Ligand spatial activity scores in 2D and 3D cellular neighborhoods

To estimate the spatial activity of a specific ligand, we first downloaded manually-curated, literature-supported receptor ligand pairs from the CellChat Human database [42] and selected those in which both the receptor and the ligand were present in our 960-gene panel. For each center cell, we quantified the spatial activity of 164 ligands in its 2D and 3D cellular neighborhoods. To quantify the activity of each ligand in a given cellular neighborhood, we first evaluated single pairs of interacting cells comprising the center cell and one of its neighbors. For each pair, we computed the geometric mean of receptor expression in the center cell and ligand expression in the neighbor cell. In this way, we required the non-zero expression of both receptor and ligand to have positive pair scores. We then compute the overall ligand activity score for a specific cellular neighborhood summing all the pair scores having the center cell as receiver. In the case of ligands paired with multiple receptors, we summarized the ligand activity as the sum of the interactions with each of the associated receptors.

### Niche enrichments

To identify the preferential localization of a group of cells *g* in a specific niche *n*, we compared the observed and expected counts of *g* cells in niche *n*. The expected counts are computed under the assumption of random, independent distribution of cells belonging to group *g* and niche *n*. To compute expected counts, we simply multiplied the total abundance of g cells in the TME and the fraction of niche n in the TME. We then compute the log2 fold changes as the log2 of the ratio between observed and expected counts. In this way, we quantified the enrichment of infiltrating tumor cells, fibroblast states and cell types across all 3D niches.

### Quantification and visualization of 3D cellular density

We quantified the 3D spatial density of a group of cells g and visualized its 2D projection using the geom_density_2d_filled function from the ggplot2 package using alpha=0.5 and h= 5.6*700. In this way, we quantified and visualized the 3D density of mesenchymal-like tumor cells (defined as pseudotime rank > 0.75), vascular endothelium and pericytes, macrophages, myofibroblasts and cytotoxic T cells.

### 3D rendering of tissue niches, gene expression and ligand activity scores

Three-dimensional visualizations were generated by downscaling and mapping the spatial coordinates of segmented cell centroids onto 300×300 images along the xy-axes, using data from sections 10, 16, 22 and 28 along the z-axis. From these, 300×300×100 voxel data was constructed by interpolating 25 intermediate sections between each of the four sections using weighted Convolutional Wasserstein barycenters [83]. Voxel data were exported as TIFF files and visualized using ParaView v5.10 as surface or volumetric representations.

### Statistical methods

Cluster-specific marker genes used for cell type and fibroblast state annotation, niche-specific differentially expressed genes in a specific cell type (including tumor cells, fibroblasts and macrophages) and niche-enriched ligands (including EMT, macrophage, dendritic, and T cell niches) were identified using a Wilcoxon Rank Sum Test comparing the expression of all genes in the group of interest versus all the remaining cells as implemented in the FindMarkers function in the Seraut R package (v. 4.0.4). p-values added to violin and boxplots were computed through an unpaired t-test using the stat_compare_means function of the ggpubr R package (v 0.6.0). Computed p values were adjusted using Bonferroni correction for multiple testing.

### Data and code availability

Image-based spatial transcriptomics are deposited on Zenodo at the following DOI 10.5281/zenodo.7899173 under restricted access for reviewing purposes. Unless requested otherwise during the review process, open access to raw and processed data will be granted upon acceptance of the manuscript, together with the release of the full analysis code through our GitHub page https://github.com/rajewsky-lab. Furthermore, it will be possible to interactively explore the analyzed data through an interactive webpage.

---

**Extended data figure 1 related to figure 1.**
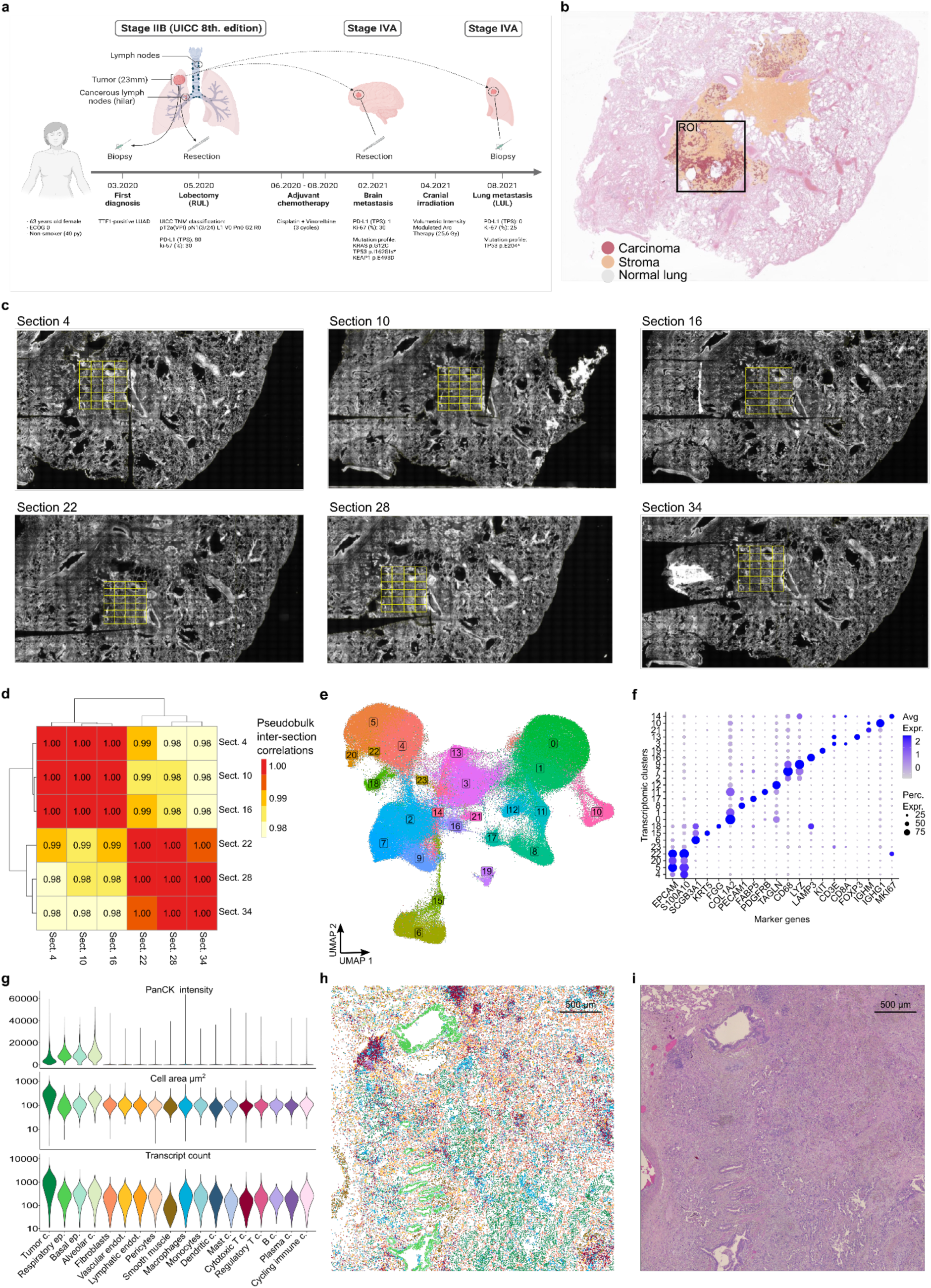
**a)** Schematics of characteristics, staging and clinical history of the NSCLC patient under study. We investigated the lobectomy sample. **b)** Deep learning-based whole slide H&E annotation of carcinoma, stroma, and normal lung regions. Black box: profiled region of interest (ROI). **c)** Whole slide images of the sections profiled with CosMx. Yellow: 24 fields of view in each ROI. **d)** Correlation of gene expression across sections. Section-to-section Pearson correlation of aggregate gene expression profiles. **e)** UMAP plot of segmented cells. Cells are colored according to their transcriptomic cluster assignment. **f)** Cluster expression of canonical markers used for cell type annotations. **g)** Cell type distributions of pancytokeratin (panCK) immunofluorescence signal (top), cell area from the segmentation masks (middle) and transcript counts (bottom). **h)** Spatial plot of cell types in section 4. **i)** Tissue morphology by conventional hematoxylin and eosin staining in section 3.

**Extended data figure 2 related to figure 2.**
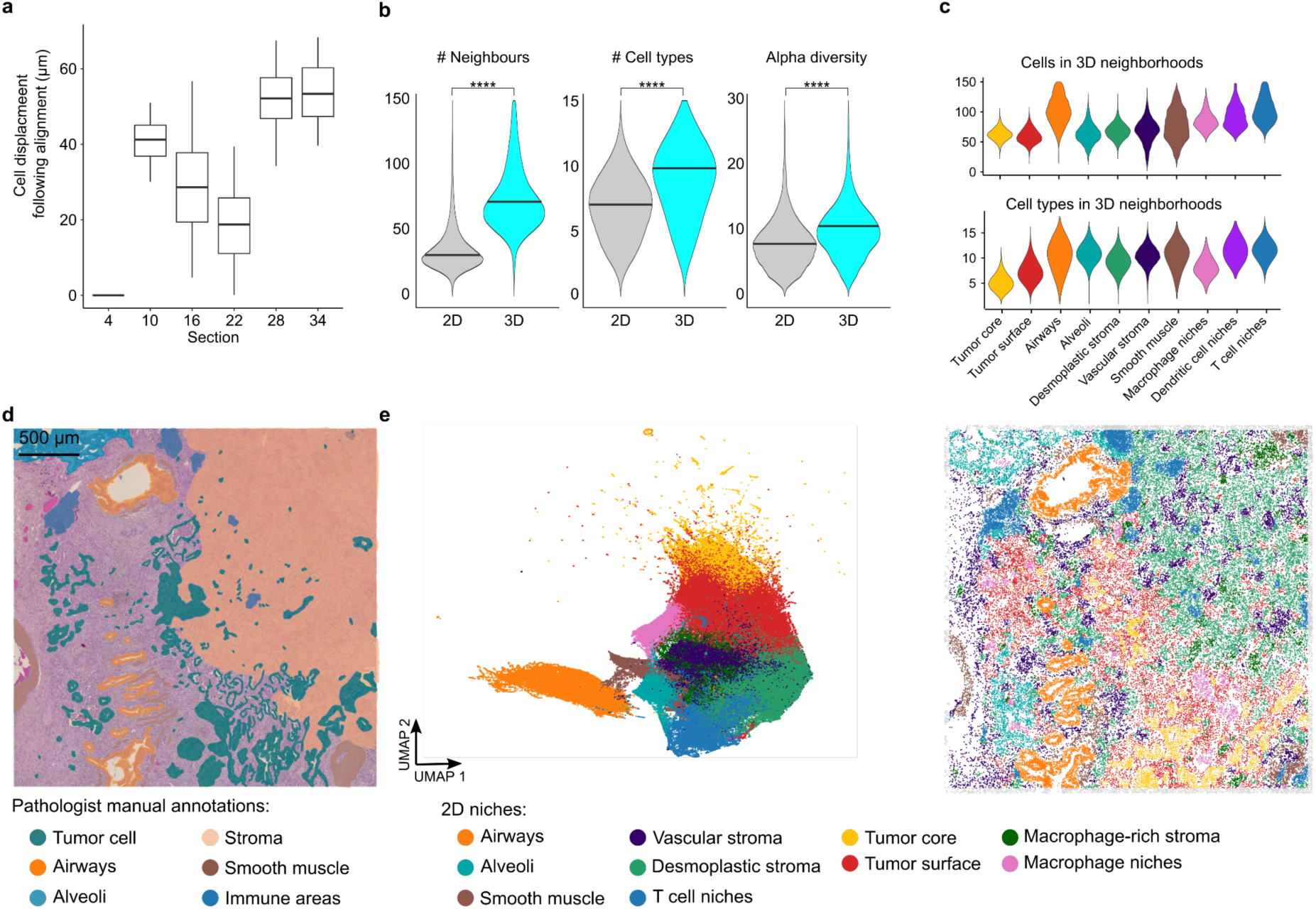
**a)** Boxplot of cellular displacement following 3D alignment. The distribution of cellular shifts for each section is shown. Section 4 was used as the anchoring section. **b)** Richness of 2D and 3D cellular neighborhoods. The distribution for the total number of neighbors, the number of different cell types and the alpha diversity (Chao index) are shown for 2D and 3D neighborhoods. ****: t-test p-values < 0.005, n= 218,378. **c)** Violin plots of the number of neighbors (top) and different cell types per 3D neighborhood for cells in different 3D niches. **d)** Pathologist manual annotation of tissue domains superimposed to hematoxylin and eosin (section 3). **e)** Left: UMAP of 2D cellular neighborhoods. Cells are grouped based on their 2D neighborhood composition and colored by 2D niche assignment. Right: Spatial map of 2D multicellular niches (section 10), gray: cells within 50 µm of the section edge.

**Extended data figure 3 related to figure 3.**
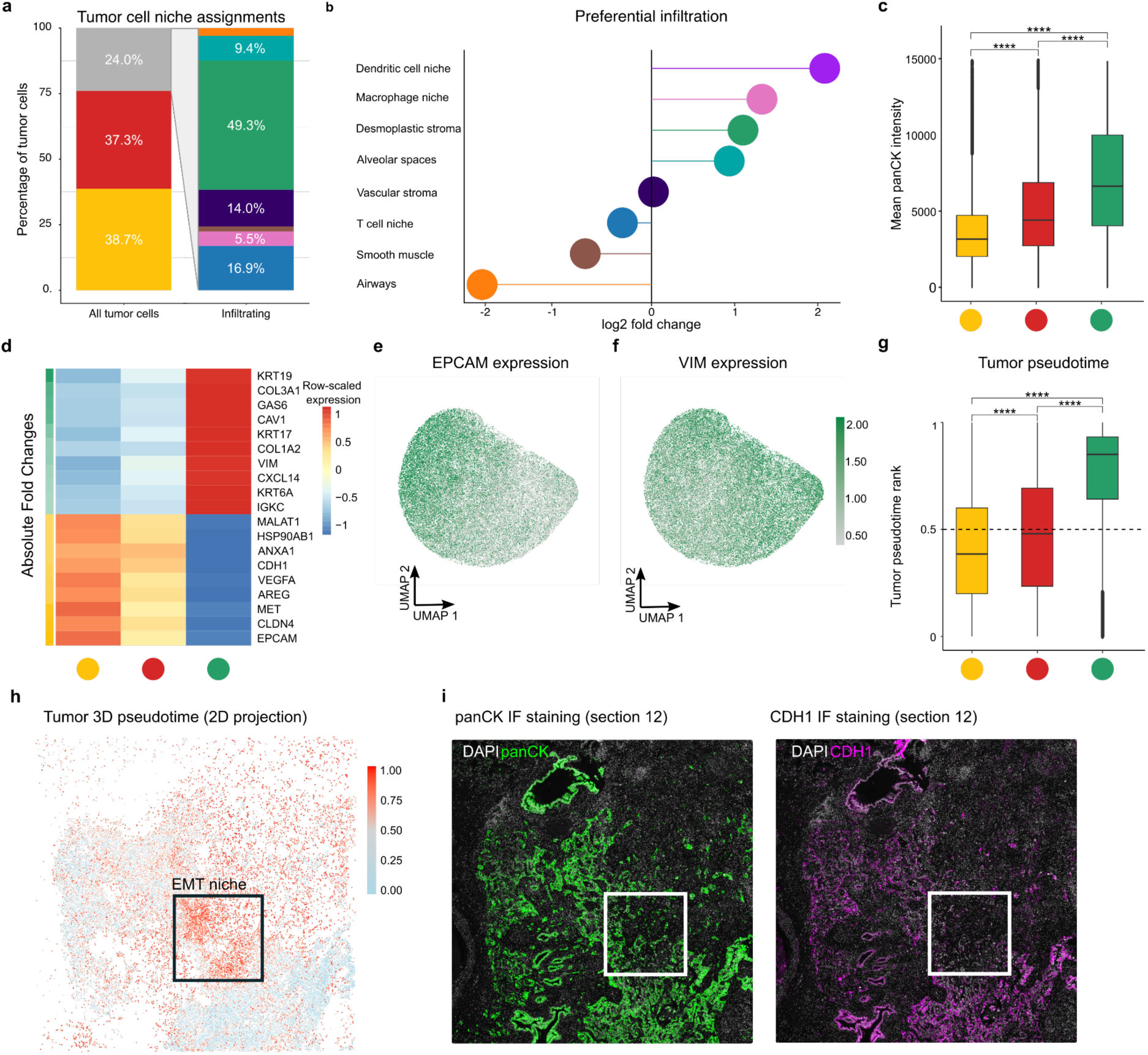
**a)** Tumor cells quantification across all niches. The left barplot includes all tumor cells (yellow: tumor core, red: tumor surface, gray: outside the tumor bed), while the right one quantifies the niche assignments of infiltrating tumor cells. Color legend in panel b. **b)** Preferential localization of infiltrating tumor cells in 3D niches. log2 fold changes are computed from the ratio of the observed vs expected number of infiltrating tumor cells per niche. **c)** Protein validation of keratin upregulation in infiltrating tumor cells. Quantification of pan-cytokeratin (panCK) immunostaining intensity in tumor cells located in the tumor core, surface and desmoplastic stroma. ****: t-test p-values < 0.005, n= 38,804. **d)** Molecular gradients at the tumor surface. Gene expression heatmap for the top differential genes between tumor cells in the tumor core and desmoplastic stroma. Row-scaled average gene expression values for tumor cells in the tumor core, surface and desmoplastic stroma. **e-f)** Epithelial to mesenchymal transition dynamics in tumor cells. Tumor expression of (e) epithelial state marker *EPCAM* (Epithelial Cell Adhesion Molecule) and (f) mesenchymal marker *VIM* (Vimentin). **g)** Pseudotime progressively increases moving from the tumor core to the tumor surface and desmoplastic stroma. Boxplot of the pseudotime rank distribution of tumor cells assigned to the tumor core, surface, and desmoplastic stroma. ****: t-test p-values < 0.005, n= 38,804. **h)** Spatial distribution of tumor pseudotime scores. **i)** Immunofluorescence validation of CDH1 downregulation in the EMT niche (section 12). Left: Immunostaining for nuclei (DAPI: white) and tumor and normal epithelial cells (panCK: green). Right: Immunostaining for nuclei (DAPI: white) and epithelial phenotype (CDH1: magenta), white square: EMT niche.

**Extended data figure 4 related to figure 4.**
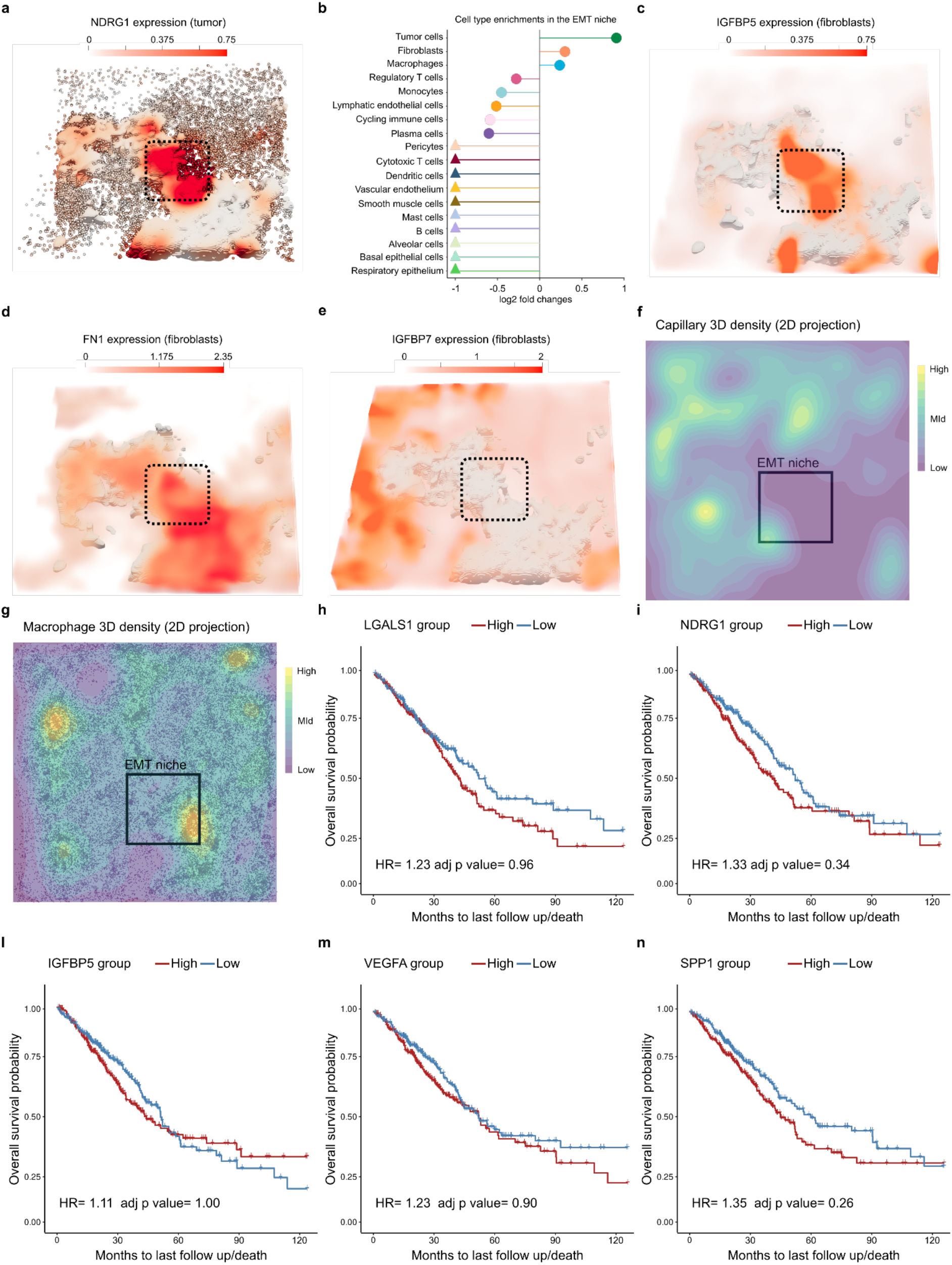
**a)** *NDRG1* expression is specific for tumor cells in the EMT niche (3D surface rendering), black box: EMT niche. **b)** Preferential localization of cell types in the EMT niche. log2 fold changes are computed from the ratio between the observed and expected number of cells per cell type in the EMT niche. **c-e)** 3D volumetric rendering of *IGFBP5* (c), *IGFBP7* (d) and FN1 (e) fibroblast expression. Gray: tumor bed (3D surface rendering), black box: EMT niche. **f-g)** 3D spatial density plot of vascular endothelium and pericytes (f) and macrophages (g). **h-n)** Survival plot of 503 lung adenocarcinoma patients from The Cancer Genome Atlas (bulk RNA-seq) stratified into high and low enrichment for the individual expression of *LGALS1* (h), *NDRG1* (i), *IGFBP5* (l), *VEGA* (m) and *SPP1* (n).

**Extended data figure 5 related to figure 5.**
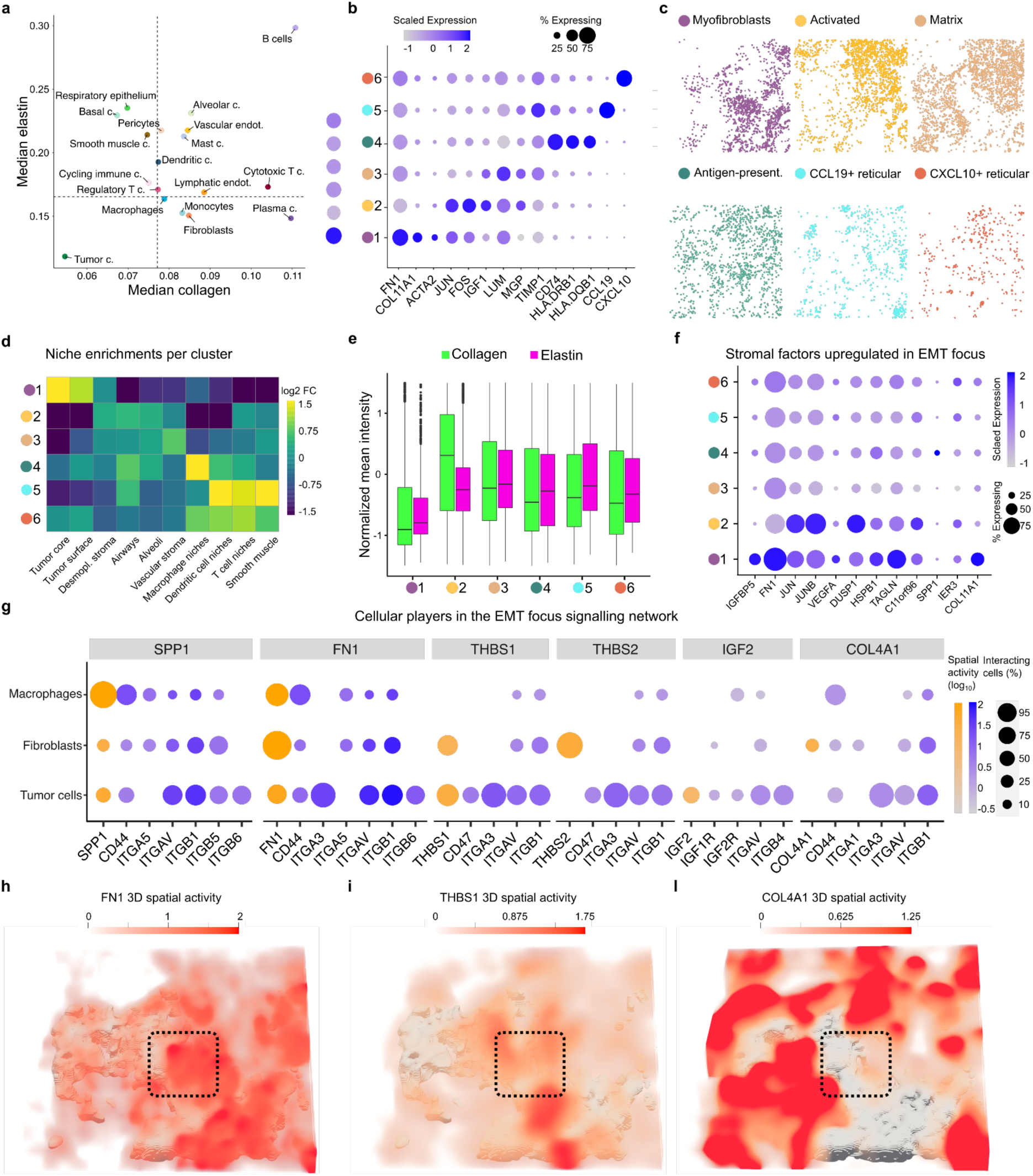
**a)** Median elastin and collagen levels surrounding cell types in section 4. Dotted lines indicate the global median elastin (horizontal) and collagen (vertical) levels. **b)** Dotplot of selected marker genes per fibroblast transcriptomic cluster. Dot size: percentage of cells expressing the gene. Dot color: scaled average expression. **c)** Spatial distribution of fibroblast clusters (section 4). **d)** Fibroblast transcriptomic states are linked with their 3D neighborhoods. Heatmap showing enrichment of fibroblast clusters in 3D niches. log2 fold changes are computed from the ratio between the observed and expected number of cells per cluster in each 3D niche. **e)** Fibroblast states are linked with ECM remodeling. Quantification of collagen and elastin (section 3) in fibroblast neighborhoods (section4). **f)** Myofibroblasts underlie the production of stromal factors specific to the EMT niche. Expression dotplot for EMT niche-specific fibroblast genes per fibroblast transcriptomic cluster. Dot size: percentage of cells expressing the gene. Dot color: scaled average expression. **g)** Cellular players of EMT niche signaling network. Dotplot of cell type-specific percentages of involved cells (dot size) and log10 average ligand sender (orange) and corresponding receptor receiver (blue) scores for ligands enriched in the EMT niche. **h-l)** FN1 (h), THBS1 (i) and COL4A1 (l) 3D spatial activity scores. Black box: EMT niche.

**Extended data figure 6 related to figure 6.**
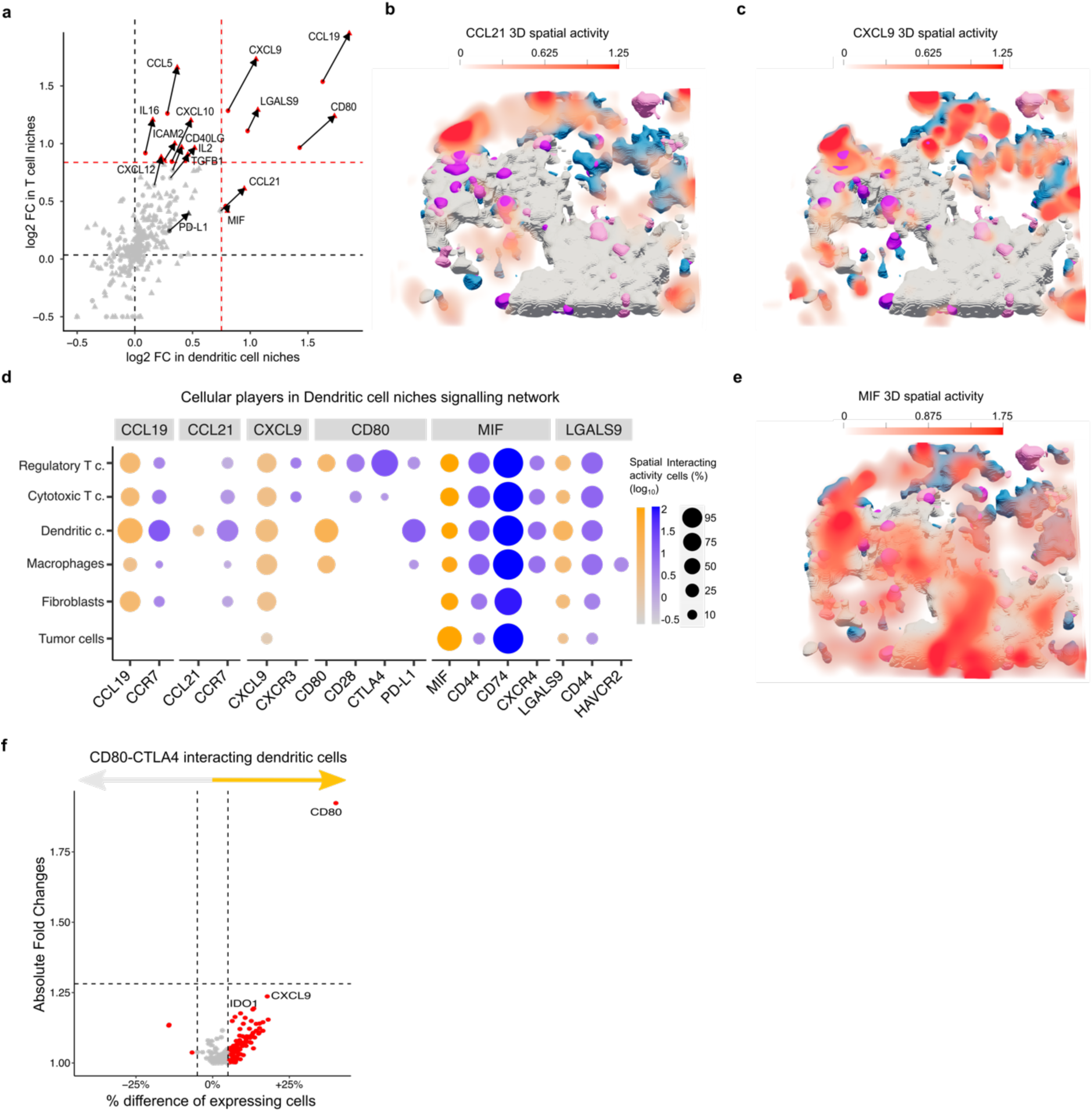
**a)** log2 Fold changes (log2 FC) for 164 ligands in ‘Dendritic cell niches’ (x axis) and T cell niches’ (y axis). FC are computed comparing average ligand sender scores for cells inside immune niches against those outside. 14 ligands with a log2FC > 0.75 in either niche (red) and PD-L1 (dark gray) are labeled. Arrows connect 2D (dots) and 3D (triangles) enrichment scores for each enriched ligand and PD-L1. **b-c)** CCL21 (b) and CXCL9 (c) 3D spatial activity scores **d)** Cellular players of Dendritic cell niche signaling network. Dotplot of cell type-specific percentages of involved cells (dot size) and log10 average ligand sender (orange) and corresponding receptor receiver (blue) scores for ligands enriched in Dendritic cell niche. **e)** MIF 3D spatial activity scores **f)** IDO1 is upregulated upon CD80-CTLA4 interaction. Differential gene expression between dendritic cells engaged in CD80-CTLA4 interaction vs those not engaged. Red: genes with more > 5% expression difference, labeled: top 3 differential genes.

