## Supplementary Material for "High-resolution molecular atlas of a lung tumor in 3D"

Tancredi Massimo Pentimalli et al.

Supplementary table 1 (separate file)

CosMx 1000-plex panel includes probes targeting 960 genes to profile cell types, states and cell-cell interactions and 20 negative control probes.

Supplementary Figure 1 related to figure 2

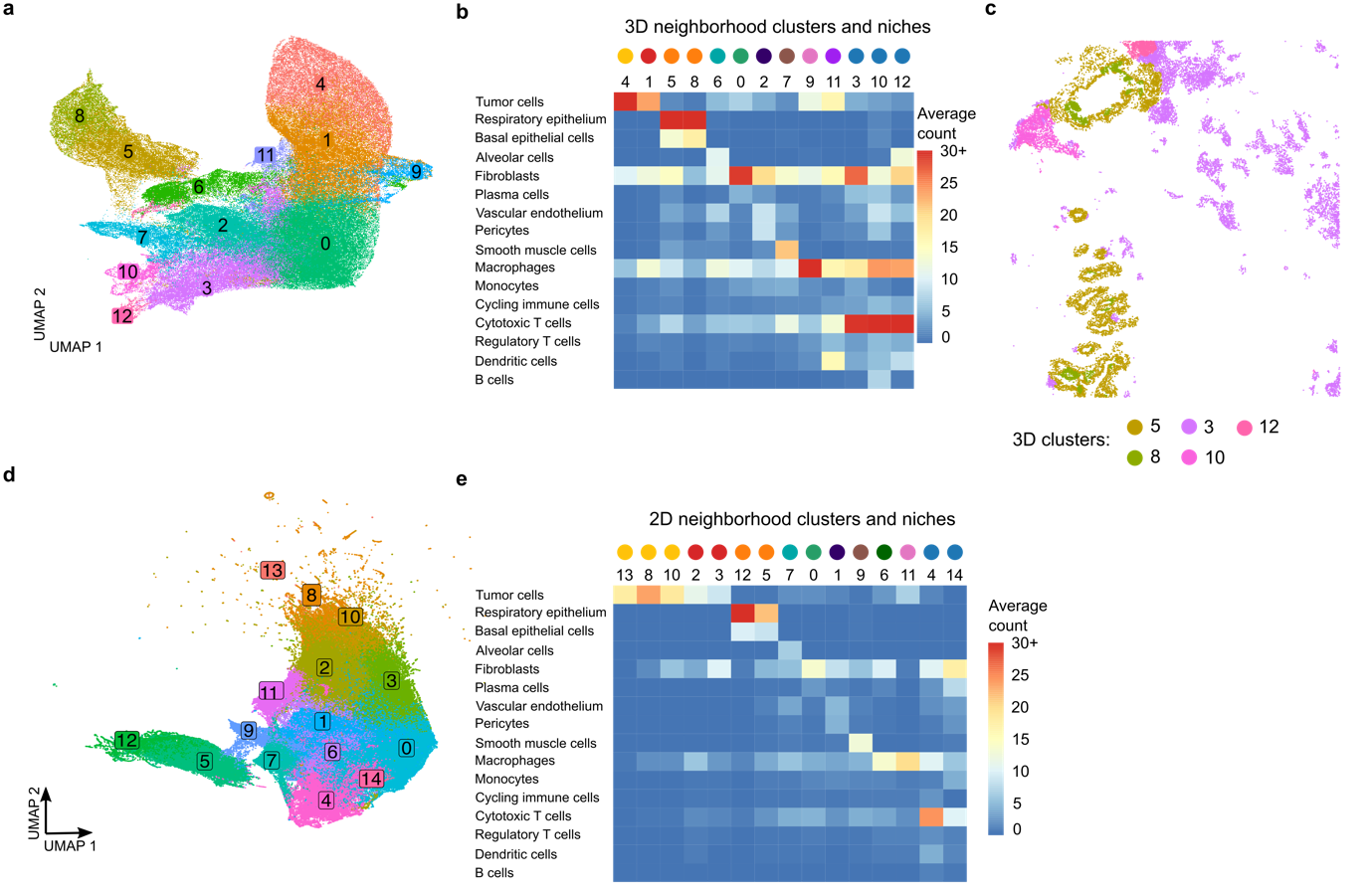

**a)** UMAP of 3D cellular neighborhoods. Cells are grouped based on their 3D neighborhood composition and colored by 3D cluster assignment.  **b)** Heatmap showing the average count of each cell type in the 3D neighborhoods of cells assigned to each cluster. Heatmap legend is clipped to 30 for visualization purposes. **c)** Clusters with similar composition and spatial pattern are merged together: Clusters 5 and 8 as ‘Airways’, Cluster 3, 10 and 12 as ‘T cell niches’. **d)** UMAP of 2D cellular neighborhoods. Cells are grouped based on their 2D neighborhood composition and colored by 2D cluster assignment. **e)** Heatmap showing the average count of each cell type in the 2D neighborhoods of cells assigned to each cluster. Heatmap legend is clipped to 30 for visualization purposes.

Supplementary video 1 related to figures 3 and 4 (separate file)

The ‘tumor surface’ covered the ‘tumor core’ and they precisely interlocked in 3D. Tumor infiltrating cells extend beyond the boundaries of the tumor surface. While their presence in alveoli, macrophage and dendritic cell niches could be explained by their spatial continuity with the tumor surface, tumor cells infiltrating the desmoplastic stroma extended away from the tumor surface. Pseudotime captured the dynamic molecular processes accompanying tumor invasion and ranked tumor cells from 0 (early) to 1 (late). Pseudotime mapped tumor cells undergoing pro-invasive epithelial-to-mesenchymal transition (EMT) not only in the desmoplastic stroma but already in one particular region of the tumor surface: the EMT niche. A multicellular signature combining tumor (*LGALS1* and *NDRG1*), fibroblast (*IGFBP5* and *VEGFA*) and macrophage-specific (*SPP1*) genes upregulated in the EMT niche robustly identified patients with shorter overall survival.

Supplementary video 2 related to figure 6 (separate file)

To identify immunomodulatory interactions underlying immune recruitment and remodeling, we focused on interactions that distinguished ‘dendritic cell niches’ and ‘macrophage niches’ from the rest of the TME. Compatible with their role in immune cell recruitment (i.e. chemotaxis), several chemokines signals (CCL19, CCL21, CXCL9) regulating the homing of dendritic and T cells in lymphoid tissues marked location of ‘dendritic cell islands’ and ‘T cell niches’ in the TME. At the same time, LGALS9-CD44 promoting regulatory T cell survival and CTLA4 which inhibits cytotoxic T cell anti-tumoral responses interacting with CD80 on dendritic cells emerged as the ligands with the strongest shared upregulation across all immune niches, highlighting their importance in promoting tumor immune escape.
